# Calcium-dependent proteolysis of the chimeric androglobin reveals altered localization of its isolated globin domain

**DOI:** 10.64898/2026.06.19.733359

**Authors:** Teng Wei Koay, Carina Osterhof, Angèle Clerc, David Hoogewijs

## Abstract

Androglobin (ADGB), a protein essential for spermatogenesis, is the most structurally unusual member of the vertebrate globin superfamily. It combines a calpain-like domain with a circularly permuted globin domain containing an embedded calmodulin-binding IQ motif, an architecture suggesting complex regulatory functions that remain poorly understood. Here, we investigated whether ADGB undergoes calcium-dependent post-translational processing, like it has been described for other calpains. ADGB underwent robust proteolytic processing upon calcium stimulation, generating several stable cleavage products following ectopic expression in mammalian cells. *In vitro* proteolysis assays demonstrated that ADGB cleavage requires cytoplasmic factor(s) and is strongly enhanced by Ca^2+^. While this process is sensitive to pan-calpain inhibition, siRNA-mediated knockdown excluded calpain 1 (CAPN1) and calpain 2 (CAPN2) as primary mediators of ADGB cleavage. In contrast, depletion of the calpain small regulatory subunit CAPNS1 markedly reduced calcium-dependent ADGB proteolysis, implicating a CAPNS1-associated calcium-responsive proteolytic pathway. Domain-mapping analyses localized the major cleavage hotspot between the N-terminal calpain-like domain and the globin-containing C-terminal region, indicating that proteolysis separates the protease-like and globin modules of the ADGB chimera. The isolated globin domain displayed enhanced interaction with calmodulin compared with full-length ADGB, whereas the extended C-terminal region impeded this interaction. Furthermore, unlike full-length ADGB, the isolated globin domain exhibited preferential localization to centrosomal structures. Collectively, these findings identify calcium-dependent proteolysis and altered subcellular localization of the isolated globin domain as previously unrecognized properties of ADGB that may be relevant to its role in ciliary biology.

## Introduction

Calcium is an essential divalent cation that plays a central role in intracellular signaling throughout eukaryotic biology. Intracellular Ca^2+^ functions predominantly as a second messenger coupling extracellular stimuli to diverse cellular responses including contraction, secretion, metabolism, differentiation, and cytoskeletal regulation [1]. The remarkable signaling versatility of Ca^2+^ arises from its flexible coordination chemistry, allowing reversible interaction with a wide variety of calcium-binding proteins and signaling effectors. Among the major calcium-responsive protein families are the calpains, a superfamily of intracellular cysteine proteases whose activity is tightly linked to calcium signaling. Calpains are Ca^2+^-dependent, non-lysosomal neutral cysteine proteases that are broadly expressed across eukaryotes. The classical calpains CAPN1 and CAPN2 are heterodimeric complexes composed of a catalytic 80-kDa large subunit associated with a common 28-kDa regulatory small subunit, CAPNS1/CAPN4 [2]. Structurally, CAPN1 and CAPN2 contain a protease core domain (CysPc domain), followed by a C2-like domain and a penta-EF-hand calcium-binding domain, whereas CAPNS1 contains a glycine-rich region and a penta-EF-hand domain involved in heterodimer stabilization [3, 4]. Calcium binding induces conformational rearrangements that promote catalytic activation of the protease core [5]. In addition to substrate cleavage, several calpains themselves undergo regulated proteolytic processing. CAPN1, CAPN2, and CAPNS1 all display stepwise cleavage events associated with activation or regulatory transitions [6–8]. Likewise, CAPN7 undergoes proteolytic removal of its N-terminal microtubule-interacting and transport (MIT) domain [9], whereas CAPN3 displays a rapid 2-step autoproteolysis for activation, as facilitated by calmodulin binding [10–13]. Collectively, these observations highlight the close functional relationship between calcium signaling and proteolytic regulation within the calpain superfamily.

Androglobin (ADGB) is a highly unusual and evolutionary distinct member of the globin superfamily that was initially classified among the non-classical calpains because of its N-terminal calpain-like domain [14–16]. In contrast to classical calpains, however, ADGB lacks the canonical penta-EF-hand calcium-binding domain. Instead, ADGB contains a circularly permuted globin domain harboring an embedded IQ motif predicted to interact with calmodulin. Recent biochemical and structural studies demonstrated that the ADGB globin domain exhibits atypical nitric oxide-related biochemical properties, including pronounced nitrite reductase activity and unusual heme coordination, suggesting that ADGB functions as a specialized signaling globin rather than a conventional oxygen transport protein [17]. Beyond canonical oxygen transport and storage, globins are increasingly recognized as versatile signaling proteins involved in nitric oxide metabolism, reactive oxygen species homeostasis, and redox-sensitive cellular signaling pathways [18]. Notably, several globins have been reported to exhibit specialized biochemical properties and restricted tissue expression patterns suggestive of regulatory rather than respiratory functions [19–21]. In this context, the unusual multidomain organization and restricted expression profile of ADGB have raised considerable interest regarding its potential role as a signaling globin within specialized cellular environments.

Interestingly, our previous overexpression studies in *Sf9* insect cells revealed the presence of multiple ADGB cleavage products, suggesting that ADGB undergoes post-translational proteolytic processing [22]. However, the proteases involved, the calcium dependency of this process, and its potential functional consequences remained unresolved. Given the presence of a calpain-like domain together with a calmodulin-binding IQ motif within ADGB, we hypothesized that calcium signaling may regulate ADGB proteolysis and potentially modulate the activity or localization of its globin domain.

In the present study, we investigated whether ADGB undergoes calcium-dependent proteolytic processing and examined how such processing influences the biochemical and cellular properties of the ADGB globin domain. We demonstrate that ADGB undergoes robust calcium-dependent cleavage mediated through a CAPNS1-associated pathway, generating stable cleavage products that separate the protease-like and globin-containing regions of the protein. Furthermore, we show that proteolytic liberation of the globin domain enhances calmodulin interaction and promotes preferential localization to centrosomal structures. Together, these findings identify regulated proteolysis as a central feature of ADGB biology and suggest a potential connection between calcium signaling, globin regulation, and ciliary function.

## Results

### ADGB is proteolyzed in a calcium-dependent manner

We previously reported ADGB to instantaneously produce several smaller products, when exogenously expressed in *Sf*9 insect cells [22], a proteolysis event that is common in calpains [9, 23]. Similarly, transient overexpression of ADGB in HEK293T cells resulted in spontaneous proteolysis into several smaller products. Interestingly, treatment of the cells with calcium in combination with a calcium ionophore further enhanced ADGB proteolysis, leading to the appearance of several distinct and specific cleaved products (**Figure 1A**). This cleavage pattern was highly robust across independently generated recombinant constructs carrying different tags, including GFP, GST, and HaloTag (**Figure 1B-D**), suggesting that ADGB proteolysis is regulated in a calcium-dependent manner and may be linked to calcium signaling pathways. We further validated this phenomenon by generating constitutively expressing cellular models, using CRISPR-based gene knock-in in HEK293T, HeLa, and MCF-7 cell lines. In all these cell types, calcium-mediated proteolysis was consistently observed (**Figure 1E-G**), indicating that ADGB cleavage under calcium signaling is a ubiquitous event that can take place in ADGB expressing cells. Since ADGB is transcriptionally silent in most cell types [24, 25], we next investigated calcium-mediated ADGB cleavage under more endogenous conditions using CRISPR activation (CRISPRa). Although CRISPRa-mediated induction produced lower overall levels of endogenous ADGB, in line with low endogenous expression levels [25], calcium treatment again resulted in the accumulation of a proportionally smaller cleavage product accompanied by a reduction in full-length ADGB levels (**Figure 1H**). These findings further support the notion that ADGB undergoes Ca^2+^-dependent proteolytic processing. In order to investigate if ADGB can be cleaved from the release of cytoplasmic Ca^2+^ alone, ADGB expressing HEK293T cells were lysed without prior calcium treatment, and subjected to further incubation, *in vitro*, at 37 °C. In the absence of EDTA, a divalent cation chelator capable of sequestering Ca^2+^, ADGB underwent rapid cleavage, generating fragment patterns similar to those observed in intact cells. Cleavage occurred immediately upon cell lysis, and the resulting fragments remained stable even after 24 hours of incubation *in vitro* (**Figure 1I**). Collectively, these findings indicate that cytoplasmic Ca^2+^ is sufficient to induce ADGB proteolysis and that this process occurs rapidly and efficiently upon Ca^2+^ exposure.

**Figure 1.**
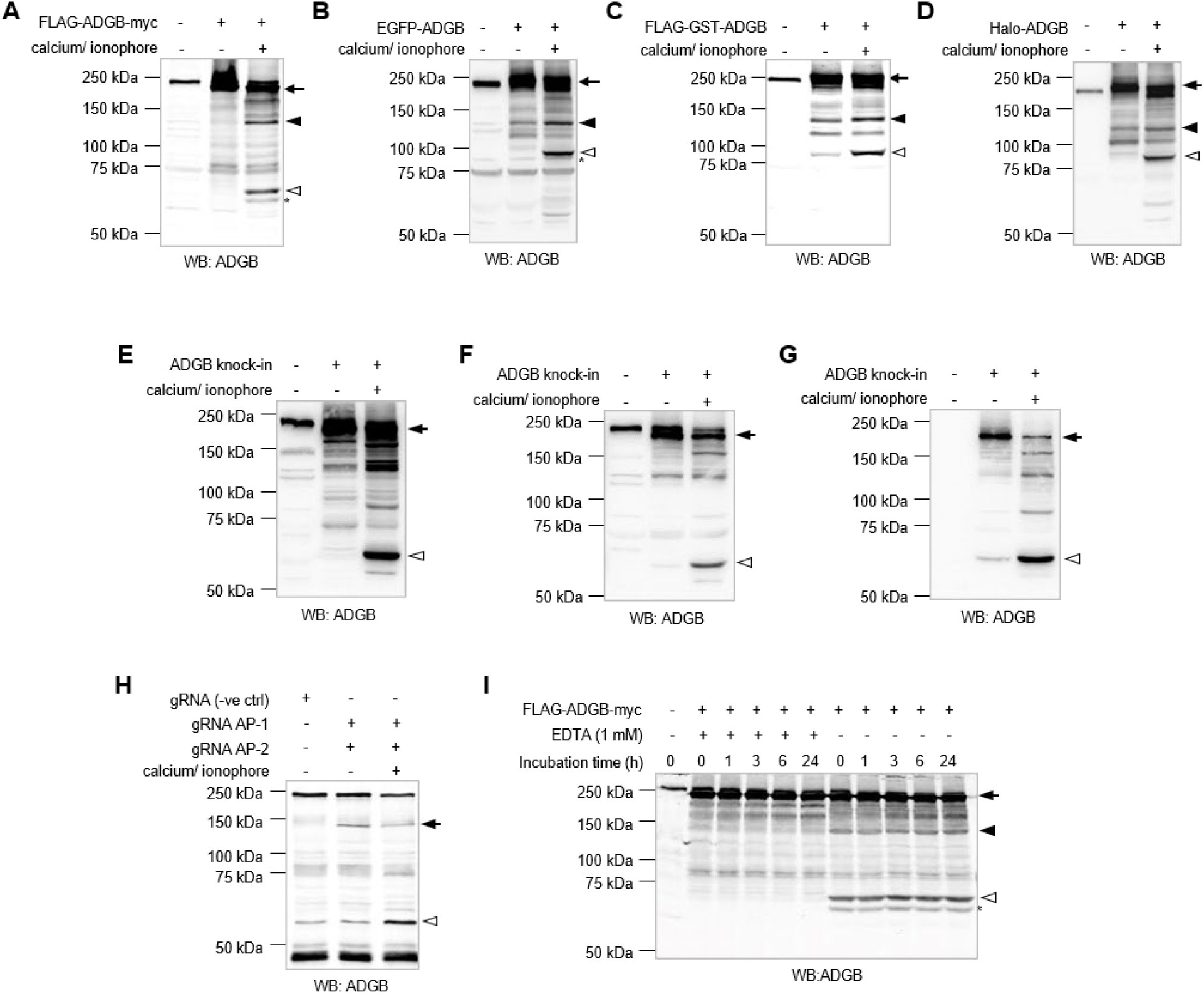
ADGB exhibits calcium-mediated proteolysis. (A-D) FLAG-ADGB-myc (A), FLAG-GST-ADGB (B), GFP-ADGB (C), and Halo-ADGB (D) were transiently expressed in HEK293T cells with and without the treatment of 50 mM CaCl_2_/ 25 μM calcium ionophore, and 100 μg of total cell lysate were analyzed by immunoblotting with an antibody against ADGB. Arrows indicate the full length ADGB with their respective protein-tag as designed in the plasmid. Open and closed triangles indicate the noticeable cleaved ADGB products that appear or increase in amount under calcium treatment. Asterisk indicates a small cleaved ADGB product that is only noticeable with the FLAG-ADGB-myc, which has smaller tags at both N- and C-termini. (E-G) Calcium-mediated ADGB cleavage is also observed in ADGB knock-in cellular models with constitutive ADGB expression. Full length ADGB is indicated by an arrow. The same cleavage pattern is observed, as indicated by open triangles in all three cell lines: HEK293T (E), HeLa (F), and MCF-7 (G). (H) CRISPRa-induced endogenous ADGB as previously described (Koay et al., 2021) shows similar enrichment of cleaved product under 50 mM CaCl_2_/ 25 μM calcium ionophore treatment albeit with a smaller molecular weight which corresponds to the smaller endogenous ADGB produced by the CRISPRa system. Arrow and open triangle indicate the full-length but shorter ADGB and its smaller cleaved product, respectively. (I) Time-course incubation (0 – 24 hours) of total lysates from HEK293T cells transiently expressing FLAG-ADGB-myc, with or without EDTA metal chelator. Sample without EDTA shows rapid cleavage of ADGB upon lysis (0 hour), and cleaved products remains stable up to 24 hours. Arrow indicates full-length ADGB, while closed triangle, open triangle and asterisk indicate cleaved ADGB products.

### Calcium-dependent cytoplasmic calpain(s) are involved in ADGB proteolysis

In the effort to further dissect the process of calcium-mediated ADGB cleavage, *in vitro* proteolysis assays were carried out using GST-purified recombinant ADGB. Under basal assay conditions, GST-ADGB remained stable and did not undergo spontaneous proteolysis (**Supp. figure 1**). However, addition of the cytoplasmic fraction derived from HEK293T cells, but not the nuclear fraction, induced robust GST-ADGB proteolysis, which was further enhanced upon calcium supplementation (**Figure 2A**). Importantly, the Ca^2+^-dependent increase in GST-ADGB proteolysis was abolished by the addition of BAPTA, a calcium-specific chelator (**Figure 2B**). These results indicate that a cytoplasmic factor is involved in ADGB proteolysis in a calcium-dependent manner. Fueled by the hypothesis that a calcium-dependent protease might be involved in ADGB cleavage, we next investigated whether members of the calpain protease family play a role in this process. Employing the *in vitro* proteolysis assays, we observed that addition of the pan-calpain inhibitor, calpeptin, can effectively inhibit the calcium-dependent ADGB cleavage in the presence of cytoplasmic proteins (**Figure 2C**). This result suggests that one or more members of the calpain superfamily might be directly or indirectly involved in the post-translational processing of ADGB under calcium stimulation. Similar results were observed when HEK293T cells transiently expressing ADGB were treated with calpeptin inhibitor, as reflected by a dose-dependent inhibition of calcium-mediated ADGB proteolysis by calpeptin (**Figure 2D**). A comparable inhibitory effect was observed upon treatment with MG-132, an established proteasome inhibitor known to additionally inhibit calpain activity (**Figure 2E**). Taken together, these results suggest a possible involvement of one or more members of the calpain family in the proteolysis of ADGB.

**Figure 2.**
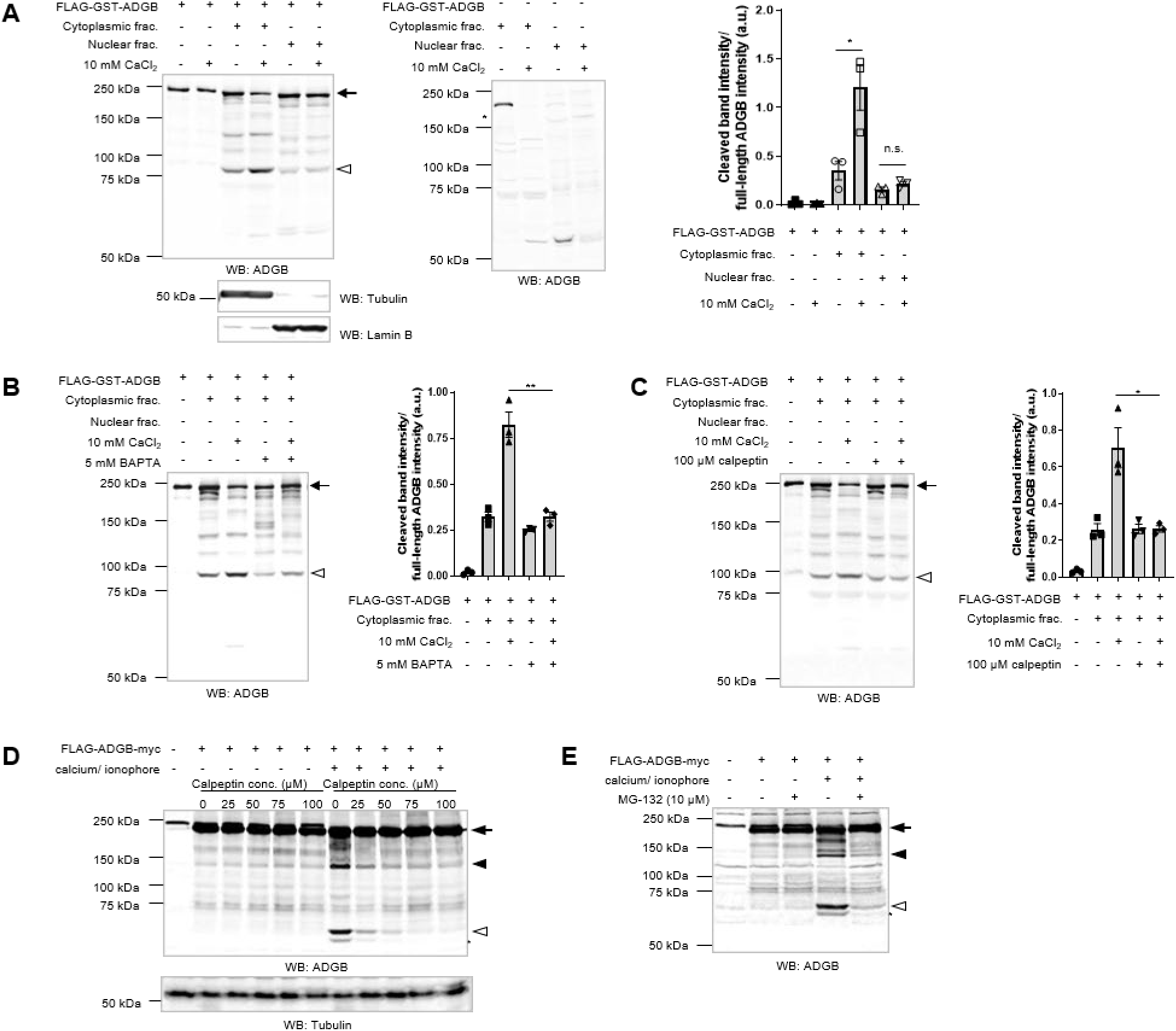
Calcium dependent cytoplasmic calpains are involved in ADGB proteolysis under calcium signaling. (A) *In vitro* proteolysis of recombinant ADGB exhibits calcium-dependent cleavage in the presence of cytoplasmic fraction. Purified FLAG-GST-ADGB (∼1.2 μg) was incubated at 37°C for 3 hours, with and without the addition of 35 μg of cytoplasmic or nuclear fraction derived from HEK293T cells, with and without the supplement of 10 mM CaCl_2_. Immunoblot against ADGB shows increased proteolysis of ADGB in cytoplasmic fraction supplemented with 10 mM CaCl_2_. Arrow and open triangle indicate full length FLAG-GST-ADGB and cleaved product, respectively. Tubulin and lamin B were used as markers for cytoplasmic fraction and nuclear fraction, respectively. A duplicated blot on the right, void of purified FLAG-GST-ADGB shows that the additional bands (marked with asterisks) are non-specific bands from the cytoplasmic or nuclear fractions. Quantification of cleaved band intensity over full-length ADGB intensity shows a marked increase in ADGB proteolysis in cytoplasmic fraction supplemented with 10 mM CaCl_2_ (N=3). (B) The addition of 5 mM BAPTA (a calcium-specific chelator) to FLAG-GST-ADGB with 35 μg of cytoplasmic fraction, supplemented with 10 mM CaCl_2_ abolished the effect of CaCl_2_ as evident from the immunoblot against ADGB and quantification of cleaved band intensity/full-length ADGB intensity (N=3). This suggests a calcium-specific activation of ADGB proteolysis in the presence of cytoplasmic proteins. (C) The addition of 100 μM calpeptin (pan-calpain inhibitor) to FLAG-GST-ADGB with 35 μg of cytoplasmic fraction, supplemented with 10 mM CaCl_2_ abolished the effect of cytoplasmic fraction/ Ca^2+^-induced proteolysis of ADGB (N=3). Quantification of cleaved band intensity/full-length ADGB intensity on the right. Arrow and open triangle indicate full-length and cleaved ADGB products, respectively. (D) FLAG-ADGB-myc was transiently expressed in HEK293T cells and treated with a titration of 25 μM, 50 μM, 75 μM, and 100 μM calpeptin for 23 hours prior to 1-hour treatment with 50 mM CaCl_2_/ 25 μM calcium ionophore. Immunoblot of cell lysate against ADGB shows a dose-dependent reduction of calcium-mediated proteolysis with calpeptin treatment. Immunoblotting with an antibody against tubulin was used as a marker for loading control. Arrow indicates full length ADGB, while closed triangle, open triangle and asterisk indicate cleaved ADGB products. (E) FLAG-ADGB-myc was transiently expressed in HEK293T cells and treated with 10 μM MG-132 (inhibitor of proteasomes, serine proteases, calpains, and other proteases) for 23 hours prior to 1-hour treatment with 50 mM CaCl_2_/ 25 μM calcium ionophore. Immunoblot of cell lysate against ADGB shows reduction in the calcium-mediated ADGB proteolysis, evident by the reduced amount of cleaved ADGB products.

### The calcium-mediated ADGB proteolysis is independent of the calmodulin-binding IQ motif on ADGB

In contrast to classical calpains, ADGB does not contain the canonical calcium-binding penta-EF hands domain. Therefore, despite our findings that ADGB proteolysis is strongly enhanced by calcium stimulation, this process is unlikely a result from direct Ca^2+^ binding to ADGB itself. However, ADGB contains a calmodulin-binding IQ motif that is flanked by the circularly permutated globin domain [14], and calmodulin binds calcium via its penta-EF hands domain. This raises the possibility that calcium might be delivered to ADGB by calmodulin via the possible interaction of calmodulin with the IQ-motif. In order to investigate this possibility, we co-overexpressed calmodulin and ADGB in HEK293T cells. However, neither cleavage of ADGB without calcium treatment, nor increase in ADGB cleavage with calcium treatment were observed (**Figure 3A**). We therefore attempted to inhibit endogenous calmodulin with W-7, a well-established synthetic antagonist of calmodulin. Similar to the overexpression experiments, W-7 treatment had no detectable effect on calcium-mediated ADGB cleavage (**Figure 3B**). These findings suggest that neither potential calmodulin -mediated calcium shuttling nor endogenous cellular calmodulin are required for ADGB proteolysis. To further validate this conclusion, we generated two ADGB variants targeting the IQ motif: an ADGB IQ-motif mutant (R918Q/R923Q) unable to bind calmodulin [26], and an ADGBΔIQ mutant lacking the entire IQ-motif. In both cases, calcium-mediated ADGB cleavage remained readily detectable (**Figure 3C,D**), further confirming that proteolytic processing of ADGB occurs independently of its calmodulin binding IQ-motif.

**Figure 3.**
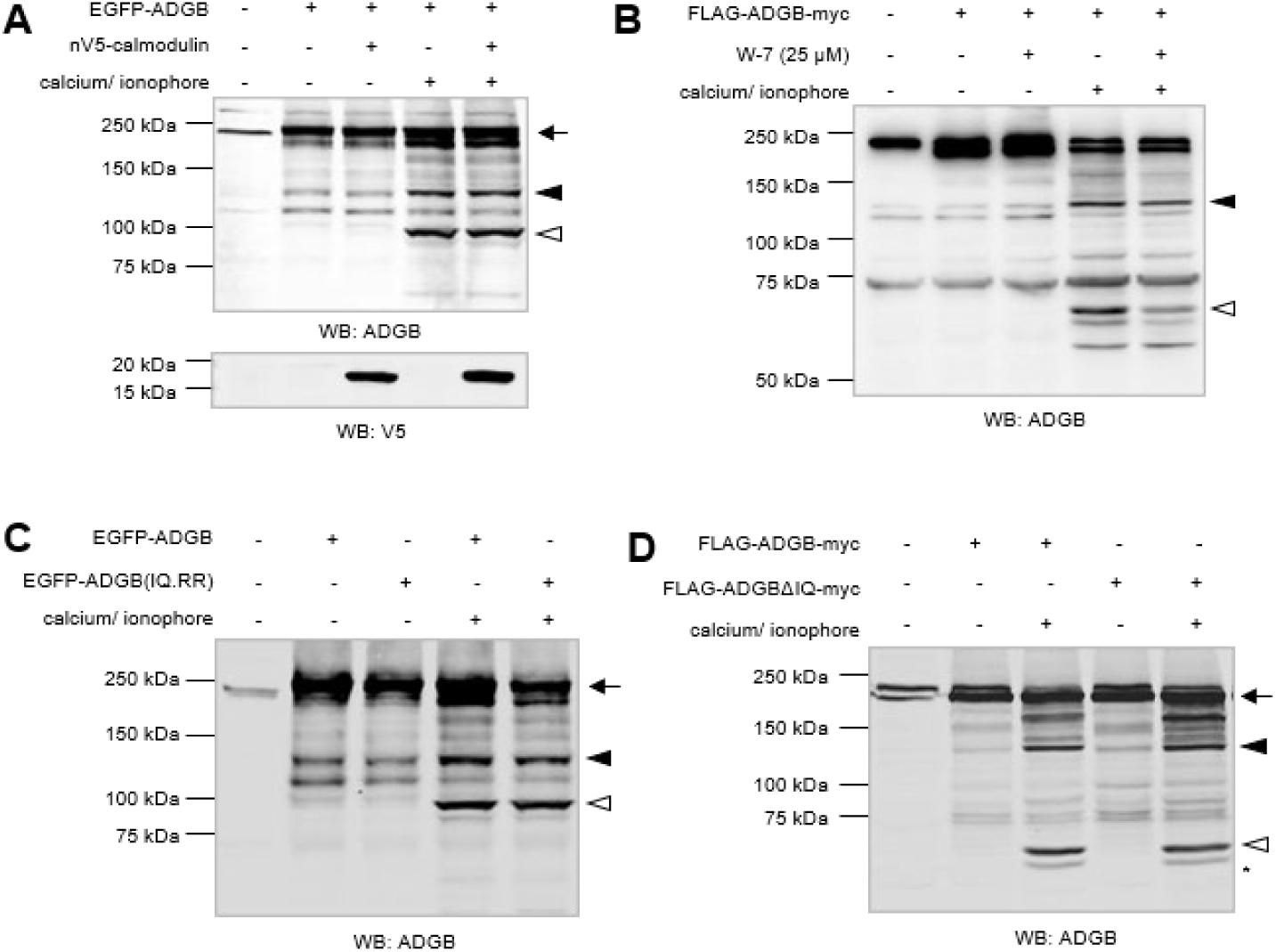
Calcium-mediated ADGB proteolysis is independent of the calmodulin-binding motif on ADGB. (A) GFP-ADGB was transiently expressed in HEK293T cells with or without co-expression of V5-calmodulin. Overexpression of V5-calmodulin does not induce the proteolysis of GFP-ADGB in the absence of calcium, and does not promote an increase in ADGB proteolysis in the presence of calcium. Immunoblotting with an antibody against V5 verifies the overexpression of V5-calmodulin. (B) Pharmacological inhibition of endogenous calmodulin with 25 μM W-7 does not prevent the calcium-induced ADGB proteolysis. Similar cleaved ADGB products were detected in the presence of calcium, with or without W-7 treatment. (C) R918Q/R923Q ADGB IQ mutant which disrupts the possible binding of calmodulin on the IQ domain (Li et al., 2013) shows no inhibition to the calcium-mediated ADGB as suggested by the similar cleaved bands that were detected in both EGFP-ADGB (wild-type) and EGFP-ADGB(IQ.RR) with calcium treatment. (D) Complete deletion of the IQ domain in ADGB does not affect the calcium-mediated proteolysis of ADGB as the cleaved products can be detected in this mutant with calcium treatment. Arrow indicates full length ADGB, while closed triangle, open triangle and asterisk indicate cleaved ADGB products.

### The calcium-dependent ADGB proteolysis is independent of the globin domain and oxygen levels

To investigate the possible role of oxygen and the ADGB globin domain on the calcium-mediated ADGB cleavage, HEK293T cells transiently expressing ADGB were cultured under normoxic (atmospheric O_2_ levels) and hypoxic (0.2% O_2_) conditions. Whereas HIF-1α protein levels increased hypoxia-dependently, we found no hypoxia-mediated changes in ADGB cleavage (**Figure 4A**). This suggests that the proteolysis of ADGB under calcium treatment is an oxygen-independent process, despite the presence of a globin domain in the ADGB chimera. At the same time, we extended our investigation to the effect of oxidative stress on this process by reoxygenation following oxygen deprivation. In contrast to decreasing HIF-1α protein levels following reoxygenation, no change was seen in the ADGB cleavage pattern (**Figure 4A**). The same was observed with ADGB deletion mutants lacking the calpain-like protease domain (FLAG-ADGBΔProtease-myc) and the calmodulin-binding IQ motif (FLAG-ADGBΔIQ-myc) (**Supp. figure 2**). Furthermore, deletion of the complete globin domain from ADGB did not affect calcium-dependent cleavage process (**Figure 4B**). Altogether, the observations indicate that modulation of oxygen levels might not be crucial at this stage of ADGB processing.

**Figure 4.**
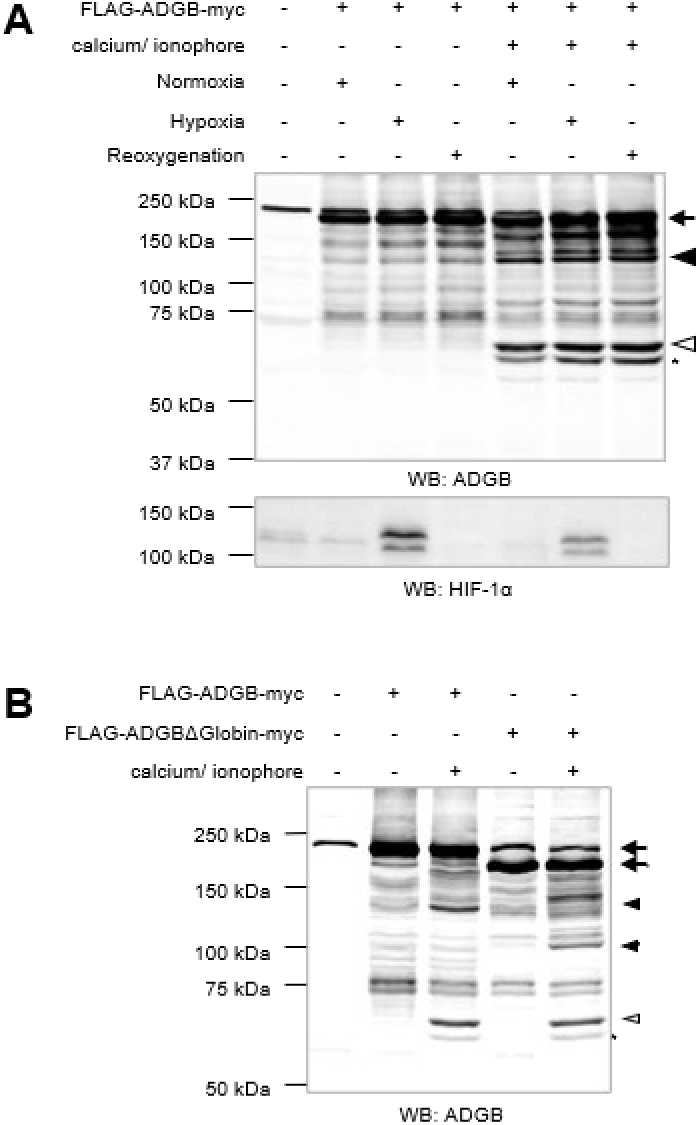
Calcium-mediated ADGB proteolysis is independent of the ADGB globin domain and oxygen levels. (A) FLAG-ADGB-myc was transiently expressed in HEK293T cells under normoxia (atmospheric O_2_ levels), hypoxia (0.2% O_2_), and hypoxia followed by reoxygenation (ROS stress). For oxygen deprivation, transfected cells were incubated under hypoxia for 24 hours and reoxygenation was carried out for 1 hour at normoxia prior to cell lysis. For samples with calcium treatment, cells were treated with 50 mM CaCl_2_/ 25 μM calcium ionophore for 1 hour prior to cell lysis. 100 µg of cell lysate were analyzed by immunoblotting with antibodies against ADGB and HIF-1α. The presence of HIF-1α in hypoxia and not in the oxygenated samples was used as a marker for oxygen deprivation. Full-length ADGB exhibits similar proteolysis with calcium treatment under normoxia, hypoxia and reoxygenation. Arrow indicates full length ADGB, while closed triangle, open triangle and asterisk indicate cleaved ADGB products. (B) Total deletion of the globin domain in ADGB shows no inhibition to the calcium-induce proteolytic event, the larger cleaved products indicated by closed triangle ([’] for mutant) shows a reduction in size proportional to the size of deletion. The smaller cleaved products, indicated by open triangle and asterisk, have the same sizes compared to that of WT.

### Calpain small subunit (CAPNS1) is involved in calcium-mediated ADGB cleavage

As we observed that the calcium-dependent ADGB proteolysis is mediated by calpain family of proteases, we next investigated whether CAPN1 and CAPN2 are involved in this process by employing the PD150606 inhibitor, with reported specificity towards CAPN1 and CAPN2 [23, 27]. In contrast to the robust inhibition observed with the pan-calpain inhibitor calpeptin, treatment with PD150606 did not affect calcium-dependent ADGB proteolysis (**Figure 5A**), suggesting that CAPN1 and CAPN2 are not the primary mediators of this cleavage event. To verify the effectiveness of PD150606, we employed mammalian-1-hybrid system with a luciferase-based readout. Briefly, presence of the PD150606 inhibitor prevents the catalytic cleavage of a flanked human spectrin (hSpec) cleavage site between a DNA-binding Gal4 domain and transcriptional activator VP16 domain, by CAPN1 and CAPN2. The intact Gal4-hSpec-VP16 regulates luciferase activity via binding and transcriptional activation of a Gal4 response element-driven promoter (**Figure 5B**). PD150606 effectively inhibited this cleavage-dependent reporter system, confirming that the inhibitor was functionally active in our assays. We next examined the role of CAPN1, CAPN2 and calpain small subunit (CAPNS1) using siRNA-mediated knockdown in HEK293T cells. Gene knock-down efficiency was verified using RT-qPCR (**Figure 5D)**. Consistent with PD150606 treatment knock-down of CAPN1 and CAPN2 had no detectable effect on the calcium-mediated ADGB cleavage. In contrast, knock-down of CAPNS1 resulted in a marked reduction in the calcium-mediated cleavage of ADGB (**Figure 5C**). Given that CAPNS1 plays a role as the regulatory subunit for both CAPN1 and CAPN2, among other functions, these findings indicate that CAPNS1 might contribute to the proteolysis of ADGB via a pathway independent of CAPN1 or CAPN2. To determine whether CAPNS1 directly interacts with ADGB, co-immunoprecipitation experiments were performed. No detectable interaction between ADGB and CAPNS1 could be observed, suggesting that CAPNS1 may not regulate this process via direct interaction (**Figure 5E**). In order to further verify the role of CAPNS1 in the proteolysis of ADGB, we performed *in vitro* proteolysis on purified recombinant GST-ADGB using cytoplasmic fractions derived from HEK293T cells subjected to CAPNS1 knock-down (**Figure 5G**), or EGFP knock-down control conditions. In agreement with the cellular experiments, cytoplasmic extracts depleted of CAPNS1 displayed a reduced capacity to mediate calcium-dependent ADGB proteolysis (**Figure 5F**). Altogether, these findings identify CAPNS1 as a regulator in calcium-mediated ADGB proteolysis.

**Figure 5.**
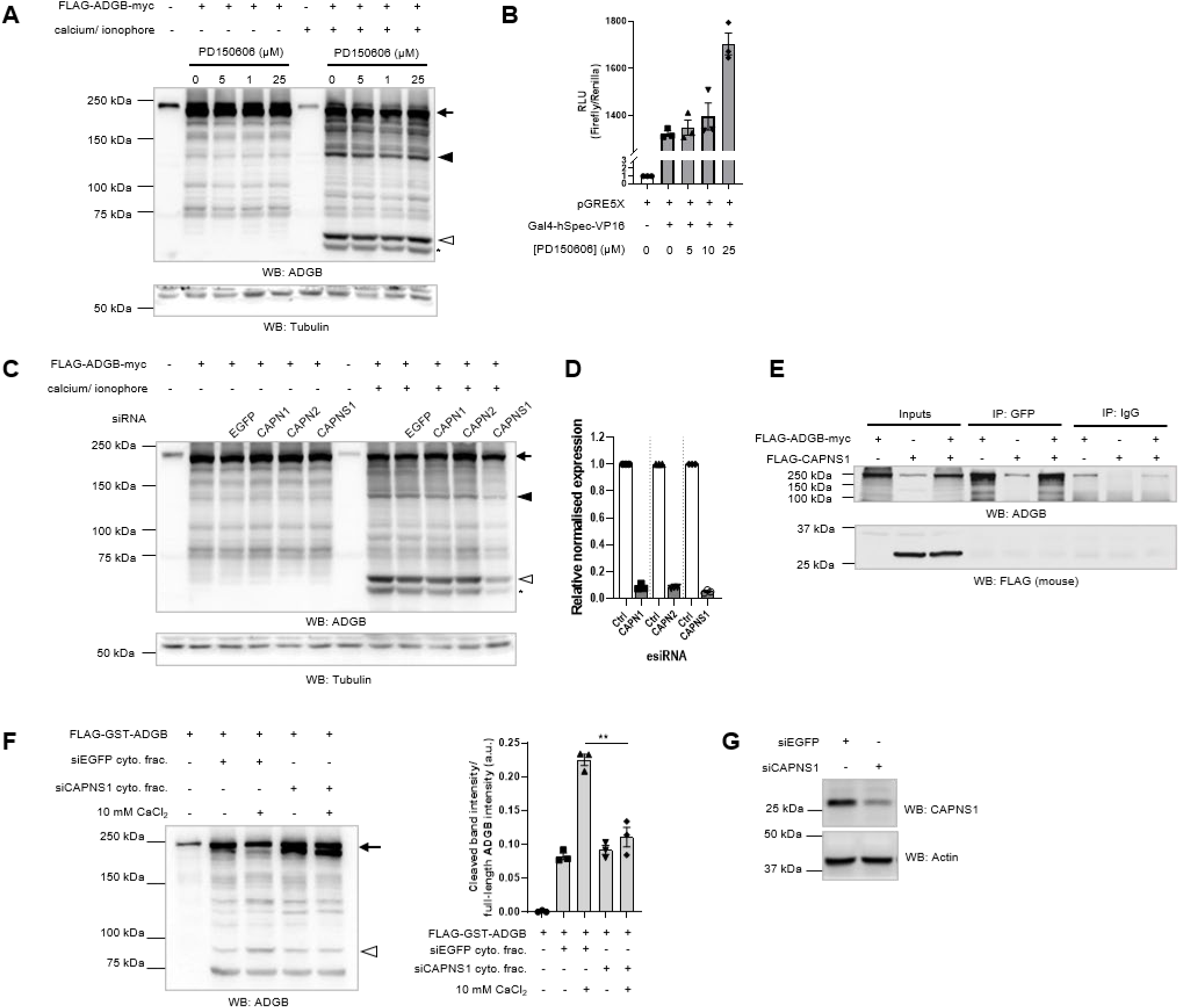
Calpain small subunit 1 (CAPNS1) is involved in calcium-mediated ADGB cleavage but not the ubiquitous calpain-1 (CAPN1) and calpain-2 (CAPN2). (A) FLAG-ADGB-myc was transiently expressed in HEK293T cells with and without 50 mM CaCl_2_/ 25 μM calcium ionophore treatment, along with increasing dosage of CAPN1- and CAPN2-specific inhibitor, PD150606 (Chukai et al., 2021; Low et al., 2014). Increasing dosage of PD150606 from 0 µM to 25 µM shows no effect on the calcium-mediated proteolysis of ADGB as displayed in anti-ADGB immunoblot. Full length ADGB is indicated by arrow, whereas cleaved products are indicated by closed triangle, open triangle, and asterisk. Loading control is shown with an immununoblot using anti-tubulin antibody. (B) The efficiency of PD150606 in inhibiting CAPN1/2 is verified using the mammalian-1-hybrid system with Gal4-hSpec-VP16 and pGRE5xE1b in a luciferase-based read-out. Active proteolytic activity of calpains cleaves human spectrin (hSpec) and separates Gal4 and VP16, leading to lower transcriptional activation of pGRE5X. Results are displayed as ratios of firefly to *Renilla* luciferase activities in relative light units (RLU) and normalized to the pGRE5xE1b promoter alone without Gal4-hSpec-VP16. Increasing dosage of PD150606 results in increasing pGRE5xE1b promoter-driven luciferase activity (N=3). (C) Knock-down of EGFP (negative control), CAPN1, CAPN2, and CAPNS1 using gene-specific siRNA pool (MISSION^®^ esiRNA, Sigma-Aldrich) shows no effect of CAPN1 and CAPN2 knock-down on calcium-dependent ADGB proteolysis, but CAPNS1 knock-down displays noticeable reduction in ADGB proteolysis, as shown in anti-ADGB immunoblot with anti-tubulin as loading control. Full length ADGB is indicated by arrow, whereas cleaved products are indicated by closed triangle, open triangle, and asterisk. (D) RT-qPCR of cDNA generated from CAPN1-, CAPN2-, and CAPNS1 knocked-down HEK293T cells shows effective knock-down of target genes. (E) Co-immunoprecipitation of FLAG-ADGB-myc and V5-CAPNS1 in HEK293T cells. Recombinant proteins were transiently expressed and cell lysates were then incubated with anti-rabbit IgG (negative control) or anti-ADGB antibody to bind and pulldown FLAG-ADGB-myc. The immunoprecipitates were immunoblotted against ADGB and FLAG with 2% cell lysates as input. Immunoblotting with anti-FLAG suggests that CAPNS1 probably has no direct interaction with ADGB. (F) *In vitro* proteolysis of recombinant ADGB exhibits reduced calcium-dependent cleavage in the siCAPNS1 cytoplasmic fraction versus siEGFP cytoplasmic fraction. Purified FLAG-GST-ADGB (∼2.4 μg) was incubated at 37°C for 3 hours, with the addition of 35 μg of cytoplasmic fraction with either CAPNS1- or EGFP- knock-down, with and without the supplement of 10 mM CaCl_2_. Representative immunoblot against ADGB and quantification of cleaved band intensity over full-length ADGB intensity show reduction in proteolysis of ADGB in CAPNS1 knocked-down cytoplasmic fraction supplemented with 10 mM CaCl_2_. Arrow and open triangle indicates full length FLAG-GST-ADGB and cleaved product, respectively. (G) Immunoblotting against CAPNS1 shows reduction in CAPNS1 in siCAPNS1 versus siEGFP cytoplasmic fractions.

### Calcium-dependent ADGB cleavage might release the globin domain from full length ADGB chimera

In order to characterize the cleavage pattern of ADGB under calcium stimulation, we analyzed proteolyzed ADGB products using N-terminally tagged constructs combined with immunoblotting against both ADGB and the respective tag proteins. Upon calcium treatment, ADGB reproducibly generated three prominent cleavage products, detectable by ADGB immunoblotting (**Figure 1A**). For clarity, the two major cleavage products are hereafter referred to as the 135-kDa fragment (closed triangle) and the 60-kDa fragment (open triangle) as shown in **Figure 1A**. Notably, the smallest band marked by an asterisk and the slightly larger but more prominent band marked by open triangle were collectively considered part of the 60-kDa fragment population due to their similar migration shifts across the experimental conditions described below.

To determine the orientation of these cleavage products within the full-length ADGB protein, large GFP or GST tags were introduced at the N-terminus of ADGB. Immunoblot analysis using anti-GFP or anti-GST antibodies demonstrated that the N-terminal tags were retained within the 60-kDa fragment, resulting in the expected proportional increase in apparent molecular weight (∼86 kDa). In contrast, the 135-kDa fragment lacked the N-terminal tags (**Figure 6A-B**). These results suggest that the 60-kDa fragment originates from the N-terminal portion of ADGB whereas the 135-kDa fragment belongs to the C-terminal part of ADGB (**Supp. figure 3**). To further define the domain composition of these fragments, a series of domain-specific deletion mutants was generated (overview provided in **Supp. figure 4**). The strategy relied on predicting size shifts in the cleavage fragments following removal of specific domains. Deletion of the calpain-like protease domain resulted in a proportional reduction in the size of the 60-kDa cleaved fragment, while the 135-kDa fragment remained unchanged (**Figure 6C-D**). Similarly, deletion of the proximal region of the 350-residue uncharacterized domain also reduced the size of 60-kDa fragment without affecting the 135-kDa fragment (**Figure 6E**). These findings indicate that both the calpain-like domain and the proximal part of the 350-residue region are present within the 60 kDa proteolysed fragment. Conversely, deletion of the distal region of the 350-residues uncharacterized domain, the globin domain, or either the proximal or distal segments of the 700-residue uncharacterized C-terminal domain, did not alter the size of 60-kDa fragment, but instead resulted in proportional reductions in size of the 135-kDa fragment, (**Figure 6F-H**, and **Figure 4B** for globin domain deletion). These results demonstrate that the region from the distal portion of the 350-residue uncharacterized domain towards the end of the ADGB protein is not present in the 60-kDa fragment and are found within the 135-kDa fragment. These findings further indicate that one or more calcium-dependent cleavage hotspots can be found within the 350-residue uncharacterized domain, a region which is situated C-terminal to the calpain-like protease domain and N-terminal to the globin domain (**Supp. figure 3**). Interestingly, transient overexpression of the 700-residue uncharacterized domain truncate also revealed calcium-dependent cleavage within this region (**Figure 6I**). By employing differential N-terminal and C-terminal tagging of this construct, we could infer the presence of at least 2 additional calcium-dependent cleavage sites within this region. Since both the 350-residue and 700-residue uncharacterized domains flank the N-terminal and C-terminal part of the globin domain respectively, the presence of calcium-dependent cleavage sites within this region suggest a possible liberation of the globin domain from the full length ADGB chimera under calcium signaling. Finally, in addition to the major cleavage events described above, we also observed a calcium-dependent processing withing the short N-terminal region of ADGB (**Supp. figure 5**). Notably, this cleavage pattern resembles the N-terminal processing reported for CAPN1 and CAPN2 [7, 8]. Intriguingly, predicted calpain cleavage sites in the ADGB protein overlap with our experimentally confirmed cuts (**Supp. figure 4**), showing a similar cleavage hotspot containing several high confidence sites around 550-650 AA [28].

**Figure 6.**
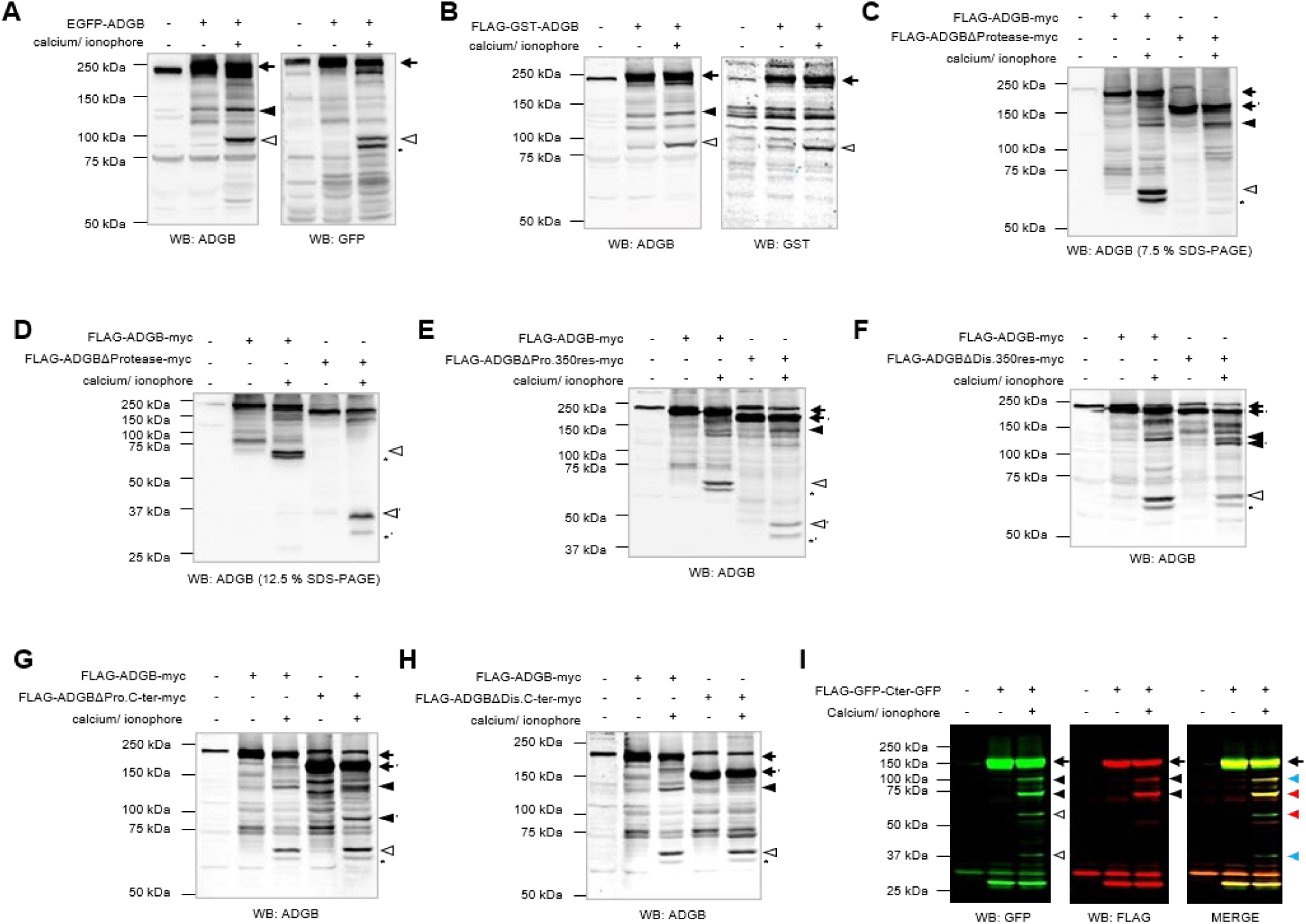
Calcium-dependent cleavage hotspots are present within N- and C-terminally situated residues in respect to ADGB globin domain. GFP-ADGB (A) and FLAG-GST-ADGB (B) was transiently expressed in HEK293T cells, with and without treatment of 50 mM CaCl_2_/ 25 μM calcium ionophore, and 100 μg of total cell lysate was analyzed with immunoblot against ADGB and GFP (A) or GST (B). For GFP-ADGB, the smaller cleaved ADGB products (with calcium treatment), indicated by open triangle and asterisk contain the N-terminus EGFP tag, whereas the larger cleaved ADGB product indicated by closed triangle, does not have the EGFP tag (A). Consistently, the smaller cleaved ADGB product of FLAG-GST-ADGB, indicated by open triangle, contains the N-terminus GST tag but not the larger cleaved ADGB product indicated by closed triangle (B). Images of immunoblotting against ADGB for GFP-ADGB and FLAG-GDT-ADGB is redundant with that of Figure 1B and 1C, respectively. (C-D) Total deletion of the calpain-like protease domain in ADGB shows a substantial reduction in size of the smaller cleaved products, indicated by open triangle and asterisk ([’] for mutant), with calcium treatment. The reduction in size is proportional to the size of deletion. Immunoblotting was performed with 7.5% (C) and 12.5% (D) SDS-PAGE to facilitate the detection of larger and smaller protein bands, respectively. Similarly, the deletion of the proximal region of the 350-residues domain (res. 410 – 588), shows a reduction in size the size of smaller cleaved products, proportional to the size of deletion (E). The smaller cleaved products indicated with open triangle and asterisk ([’] for mutant). (F-H) Partial removal of the distal region of the 350-residues domain (res. 641 – 762) (F), or the proximal region of the uncharacterized C-terminal domain (res. 988 – 1295) (G), or the distal region of the uncharacterized C-terminal domain (res. 1296 – 1667) (H), transiently expressed in HEK293T cells with and without calcium treatment. Contrary to (C-E) and (F-H), the smaller cleaved products, indicated by open triangle and asterisk ([’] for mutant), have the same sizes compared to WT, whereas the larger cleaved product indicated by closed triangle ([’] for mutant) shows a reduction in size due to the deletion. (I) ADGB C-terminal domain flanked with FLAG-GFP at the N-terminus and GFP at the C-terminus were overexpressed in HEK293T cells with and without calcium treatment. Immunoblotting analysis against GFP and FLAG shows calcium dependent proteolysis in this domain. Dual-mode imaging of GFP and FLAG was used to differentiate the N-terminal fragments (with FLAG tag) and the C-terminal fragment (without FLAG tag). Full length protein is indicated by arrow. The two bands indicated by closed arrow in FLAG WB are of the N-terminal of the truncate, whereas the two bands indicated by open triangle in the GFP WB (void of FLAG tag) made up the C-terminal of the truncate. Based on the molecular weight, it is predicted that the pairs indicated in red or blue, in the merged image, made up the full-length of the ADGB C-terminal truncate, suggesting that at least two calcium-dependent cleavage hotspots are present.

### ADGB globin domain interacts with calmodulin and this interaction is impeded by the C-terminal domain

The presence of an embedded IQ motif within the circularly permuted globin domain prompted us previously to investigate the potential interaction of ADGB with CaM [29]. In our earlier study, however, full length ADGB did not exhibit a readily detectable interaction with calmodulin. Consistent with these former observations, co-immunoprecipitation of the isolated ADGB globin domain however with calmodulin, demonstrated robust interaction between these two proteins (**Figure 7A**). This result indicates that the globin domain within the ADGB chimera might not be able to interact with calmodulin despite its IQ-motif, whereas the isolated globin domain can bind to calmodulin. To further explore this hypothesis, we generated additional fusion constructs. Co-immunoprecipitation experiments using an ADGB construct comprising the N-terminal part of the protein together with the globin domain (residue 1 to 968) still demonstrated robust interaction with calmodulin (**Figure 7B**). This suggests that the presence of the region N-terminal to the globin domain does not interfere with calmodulin binding to the IQ-motif. In contrast, an ADGB construct consisting of the globin domain together with the extended C-terminal region (700-residue uncharacterized domain) failed to efficiently interact with calmodulin (**Figure 7C**). These observations raise a possible mechanistic interference of the extended C-terminal domain on the binding of calmodulin to the IQ-motif within the globin domain, and suggest that proteolytic separation may have putative functional consequences. Consistent with this interpretation, a C-terminally GFP-tagged isolated ADGB globin domain, in contrast to its N-terminally tagged counterpart, also failed to efficiently interact with calmodulin (**Supp. figure 6**). Altogether, these experiments indicate that the ADGB globin domain, when isolated from the surrounding domains or relieved from C-terminal structural constraints, can interact more effectively with calmodulin.

**Figure 7.**
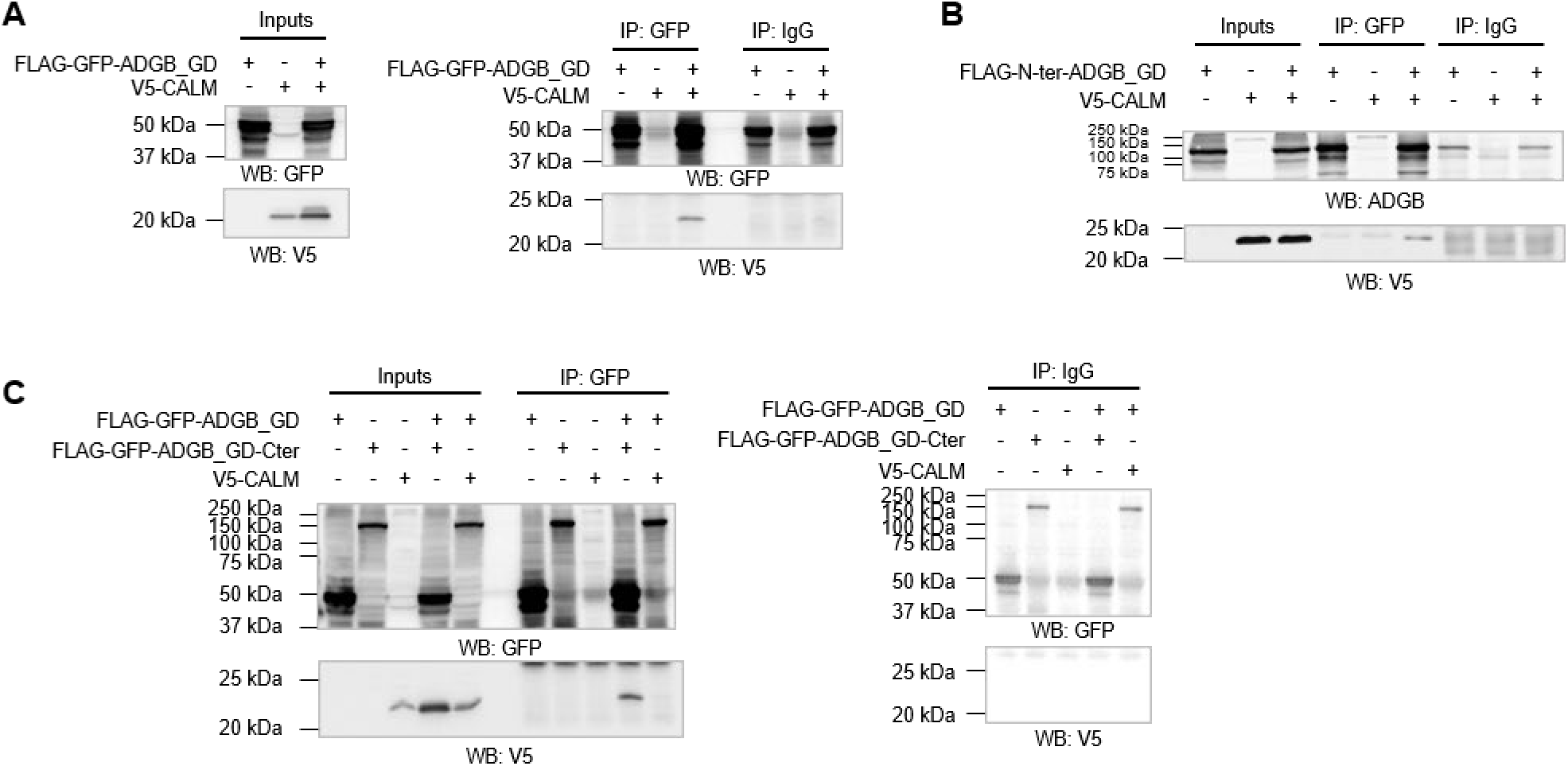
IQ-motif-containing ADGB globin domain (ADGB_GD) interacts with calmodulin and the interaction is interfered by the long C-terminal domain. (A) Co-immunoprecipitation of FLAG-GFP-ADGB_GD and V5-CALM in HEK293T cells. Recombinant proteins were transiently expressed, and cell lysates were then incubated with anti-rabbit IgG (negative control) or anti-GFP antibody to bind and pulldown GFP-tag on FLAG-GFP-ADGB_GD. The immunoprecipitates were immunoblotted against GFP and V5 with 2% cell lysates as input. V5-CALM was immunoprecipitated with FLAG-GFP-ADGB_GD as shown in the V5 immunoblot. (B) Co-immunoprecipitation of FLAG-Nter-ADGB_GD (without ADGB C-terminal domain) and V5-CALM in HEK293T cells, with protocol as mentioned in (A). Immunoblotting against V5 shows no interference in the binding of ADGB_GD in the presence of N-terminal segment and the absence of C-terminal segment. (C) Co-immunoprecipitation of FLAG-GFP-ADGB_GD or FLAG-GFP-ADGB_GD-Cter (without N-terminal segment in respective to the globin domain) and V5-CALM in HEK293T cells. The immunoprecipitates indicate that the presence of C-terminal domain (in respective to the globin domain) interferes with the binding of V5-CALM onto the ADGB_GD. Immunoprecipitation of FLAG-GFP-ADGB_GD and V5-CALM acts as a positive control.

### The ADGB globin domain preferentially localizes to centrosomal structures

To investigate the significance of ADGB proteolysis in a cellular context, we examined the subcellular localization of different ADGB truncation constructs in HEK293T cells by visualizing recombinant GFP-tagged fusion proteins. We found that the full-length ADGB (GFP-ADGB) displayed a diffuse cytosolic localization pattern throughout the cell (**Figure 8A**). A similar localization pattern was observed for the ADGB construct consisting of the globin domain together with the long C-terminal domain (GFP-ADGB_GD-Cter, **Figure 8B**). Intriguingly, ADGB globin domain alone without the long C-terminal (GFP-ADGB_GD, **Figure 8C**) displayed a distinct dot-like localization, predominantly detected in the perinuclear region. To further characterize these punctate structures, cells were co-stained with the centriole marker PCM-1. Immunofluorescence analysis revealed overlaying signals between PCM-1 and the GFP-ADGB_GD puncta (**Figure 8C**). In contrast, centrosomal localization was only rarely observed for either the full-length ADGB protein or the GFP-ADGB_GD-Cter construct **(Figure 8A-B)**. Quantitative analysis comparing centrosomal versus diffuse cytoplasmic localization further demonstrated preferential localization of GFP-ADGB_GD to the centrosome (50.6%) in comparison with the full-length GFP-ADGB and the GFP-ADGB_GD-Cter (**Figure 8D**). Mutations affecting heme-binding proximal F8 and/or distal E7 residues within the ADGB globin domain did not alter the preferential centrosomal localization, suggesting that this differential localization occurs independently of heme-binding (**Supp. figure 7).** To further explore a potential centrosomal association of the ADGB globin domain, we examined endogenous proteins identified by GFP-ADGB_GD co-immunoprecipitation and mass spectrometry [29]. Proteins exhibiting log2 enrichment ≥ 1 were analyzed for reported centrosomal or basal body localization using Gene Ontology annotations combined with subsequent Human Protein Atlas-based localization data. The enrichment of 52 endogenous centrosome-associated proteins among GFP-ADGB_GD interactors corroborates the observed centrosomal localization of the isolated globin domain (**Supp. table 1**).

**Figure 8.**
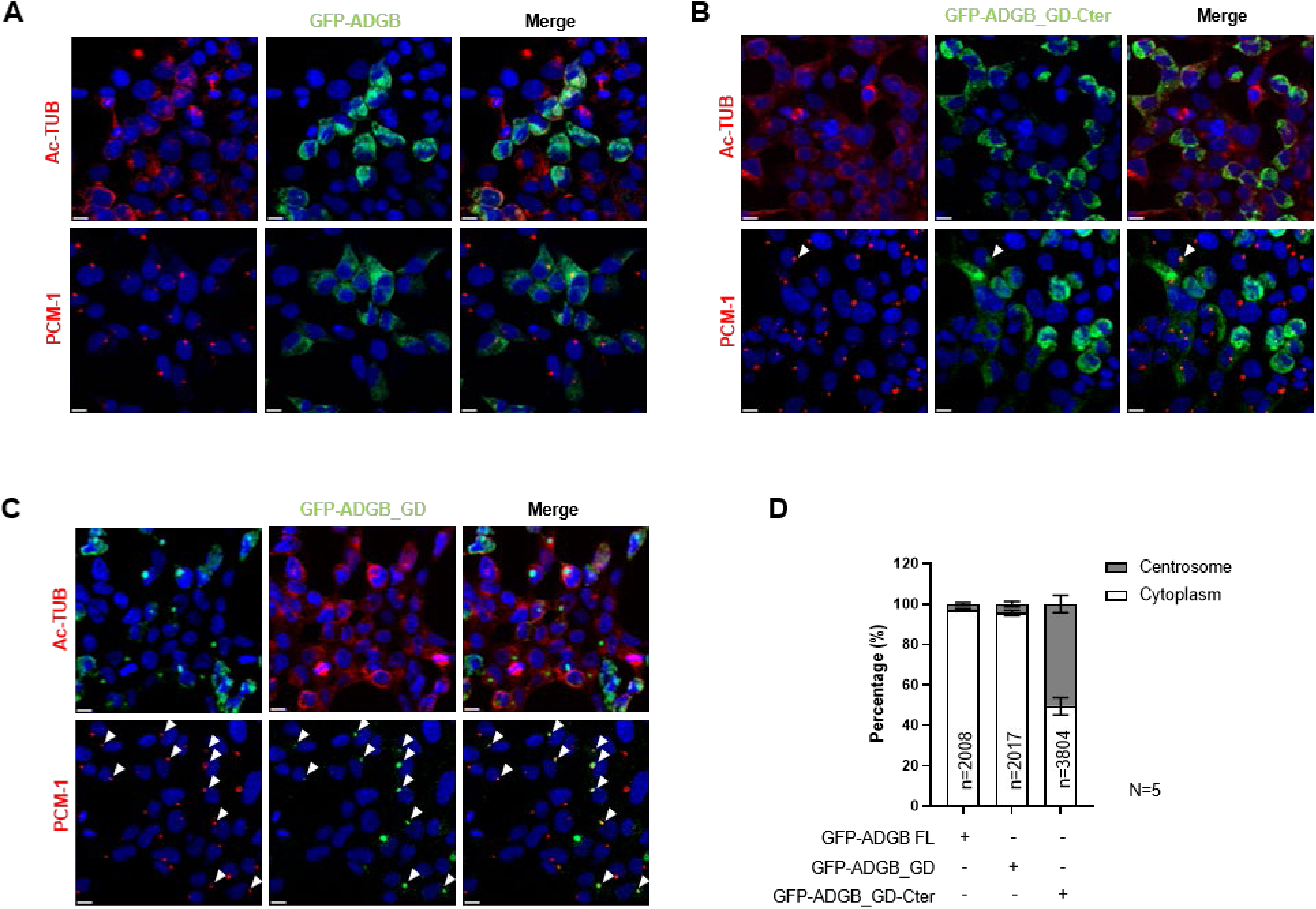
Recombinant ADGB globin domain preferentially localizes to the centrioles. (A-C) N-terminally GFP-tagged ADGB FL (GFP-ADGB) (A), ADGB globin domain with C-terminal domain (GFP-ADGB_GD-Cter) (B), and ADGB globin domain (GFP-ADGB_GD) (C) were transiently expressed in HEK293T cells for 24 hours prior to 3.7% PFA/PBS fixation. Fixed cells were stained against acetylated tubulin (Ac-TUB) or centriolar marker (PCM-1) prior to mounting with DAPI staining. Red signals are the stained Ac-TUB or PCM-1 as indicated, whereas GFP signals show the subcellular localization of ADGB FL and its truncates. GFP-ADGB and GFP-ADGB_GD-Cter are distributed in the cytoplasm, whereas GFP-ADGB_GD shows preferential localization in the centrioles (or centrosome), as evident by the overlaying with PCM-1 signal (C). White triangles show the colocalization of GFP signals to the centrosomes. Scale bar represents 10 μM. (D) Statistical analysis of the number of cells displaying localization at the cytoplasm versus centrosome. Bar chart represents the average percentages of transfected cells with GFP signals localized at the centrosome versus cytoplasm. Average percentages of centrosomally localized GFP-ADGB FL, GFP-ADGB_GD-Cter, and GFP-ADGB_GD are 2.9%, 4.5%, and 50.6%, respectively (n=5 independent experiments).

Collectively, these findings suggest that ADGB proteolysis may liberate the globin domain from the full-length chimeric protein, thereby enabling its relocalization to the centriole. Given that centrioles serve as precursors of basal bodies during ciliogenesis, these observations suggest a potential functional connection between ADGB proteolytic processing and ciliary biology.

## Discussion

Post-translational proteolysis represents a fundamental and irreversible protein modification that is often underappreciated and interpreted as protein degradation or turnover. However, limited proteolysis generating stable and functional products is undeniably physiologically relevant [30]. Classic examples of cleavage-based enzyme activations include the cleavage of Factor XIII and fibrinogen during blood coagulation [31] and the cleavage of chymotrypsinogen to chymotrypsin in digestion [32]. In this study, we report that the chimeric protein ADGB undergoes a limited proteolysis upon treatment with calcium, generating two prominent cleavage products which we termed 60-kDa fragment and 135-kDa fragment for simplicity. Size-based predictions suggest that the 60-kDa and 135-kDa fragments (∼195 kDa) are most probably the products from a single cleavage that separates the full-length ADGB. We comprehensively investigated and showed that the cleavage hotspot is situated between the calpain-like protease domain and the globin domain, with the 135-kDa fragment retaining both the globin domain and the extended C-terminal domain (700-residue uncharacterized domain). The remarkable stability and reproducibility of these cleavage products strongly suggest that ADGB proteolysis represents a regulated post-translational event rather than nonspecific degradation.

The atypical multidomain architecture of ADGB makes this observation particularly intriguing. ADGB contains an N-terminal calpain-like domain fused to a circularly permuted globin domain harboring an embedded IQ motif, followed by an extended C-terminal region of unknown function [14]. While ADGB was initially annotated as CAPN16 or “demi-calpain” [16], the absence of critical catalytic residues within the CysPc domain has long raised uncertainty regarding its intrinsic proteolytic activity. Our findings instead support a model in which ADGB itself is processed by external calcium-dependent proteases. Several observations are consistent with this interpretation. First, purified recombinant ADGB remained stable *in vitro* unless cytoplasmic fractions were added. Second, cleavage was strongly enhanced by calcium and blocked by calcium chelation. Third, pharmacological inhibition of calpain activity robustly reduced ADGB processing. Together, these findings indicate that ADGB proteolysis is mediated by an external, calcium-dependent cytoplasmic proteolytic system.

Interestingly, our data further identify CAPNS1 as an important contributor to ADGB cleavage. CAPNS1 is best known as the small regulatory subunit of the classical calpains CAPN1 and CAPN2 [33]. Surprisingly, however, neither inhibition nor knockdown of CAPN1 or CAPN2 affected ADGB processing, whereas depletion of CAPNS1 consistently reduced calcium- mediated cleavage. Although we were unable to detect a stable interaction between CAPNS1 and ADGB under our specific experimental conditions, these findings suggest that CAPNS1 contributes to ADGB processing through a non-canonical pathway independent of classical CAPN1/CAPN2 activity. Such a model is compatible with previous reports demonstrating broader signaling and scaffolding functions of CAPNS1 beyond its established role within calpain heterodimers [34–36]. Moreover, reciprocal proteolytic regulation among calpain family members has increasingly emerged as an important regulatory principle within the calpain system itself [37]. Of note, a lack of detectable CAPNS1 in our co-immunoprecipitation assays and M2H experiments (data not shown) might also reflect the transient nature of enzyme-substrate interactions, as has been described for other calpain substrates [38]. Approaches that stabilize such complexes, such as calpain inhibitor pre-treatment or catalytically inactive calpain mutants, may be required to detect and could still reveal a potential direct interaction [39].

Our data further suggests that ADGB proteolysis may be an evolutionarily conserved feature of the protein. A recent study in *Tetrahymena thermophila* expressing C-terminally tagged Adgb detected a prominent ∼130-kDa fragment from a 180-kDa full-length protein [40], closely matching the 135-kDa C-terminal cleavage product we identify in the current study. Because the *Tetrahymena* protein lacks a globin domain, this suggests that proteolytic cleavage of ADGB predates the evolutionary acquisition of the globin domain [41], consistent with our finding that the globin domain is dispensable for cleavage.

A substantial observation of the current study is that proteolytic liberation of the globin domain profoundly alters its cellular properties. Whereas full-length ADGB displayed predominantly diffuse cytoplasmic localization, the isolated globin domain exhibited striking preferential localization to the centrosome/centriole compartment. Similarly, the isolated globin domain displayed enhanced interaction with calmodulin compared with the intact protein. These findings suggest that proteolytic processing modifies the structural accessibility or conformational flexibility of the globin module. Such changes are particularly relevant given the increasing appreciation that globin function extends far beyond canonical oxygen transport and is often strongly influenced by conformational dynamics and protein-protein interactions [20].

Recent structural and biochemical studies considerably expanded our understanding of the unusual ADGB globin domain. Reeder and colleagues demonstrated that the circularly permuted ADGB globin exhibits highly atypical biochemical properties, including unusual heme coordination, pronounced nitrite reductase activity, and formation of five-coordinate NO-heme complexes more characteristic of signaling globins than classical oxygen carriers [17]. Structural analyses further revealed unique features of the ADGB globin fold, including stabilization through intramolecular disulfide interactions and replacement of the otherwise highly conserved CD1 phenylalanine residue. Together, these findings strongly support the concept that ADGB functions as a specialized signaling globin rather than a conventional respiratory globin.

Also relevant to the current study, Nie et al. recently demonstrated that calmodulin directly interacts with the ADGB globin domain through the embedded IQ motif and enhances its nitrite reductase activity [42]. These observations integrate well with our findings. While the IQ motif proved dispensable for calcium-mediated proteolytic cleavage itself, the liberated globin domain displayed enhanced calmodulin interaction and preferential centrosomal localization. One possibility is therefore that calcium-dependent proteolysis represents an upstream regulatory step that exposes or activates distinct biochemical properties of the globin module. In this context, cleavage may dynamically modulate globin accessibility, interaction networks, or signaling capacity within specialized cellular compartments.

The preferential centrosomal localization of the isolated globin domain is particularly intriguing in light of the established association of ADGB with motile ciliogenesis. We previously demonstrated that ADGB is transcriptionally regulated by FOXJ1 and RFX2, master regulators of motile ciliogenesis [43]. Furthermore, ADGB-deficient mice develop severe defects in spermatogenesis characterized by abnormal sperm head morphology, impaired manchette organization, defective acrosome formation, and complete male infertility, establishing ADGB as an essential factor in specialized flagellated cells [29]. The current findings extend this concept by suggesting that calcium-dependent proteolysis may dynamically regulate ADGB localization or function within ciliary and centrosomal compartments. Consistent with this interpretation, ADGB homologs were identified in *Tetrahymena thermophila* within the central apparatus, a structure critically involved in regulation of dynein activity and ciliary beating in motile cilia [40]. More recently, high-resolution structural and proteomic analyses of human motile cilia further localized ADGB to the C1b projection of the central apparatus [44]. These observations further strengthen the view that ADGB participates in highly specialized calcium-responsive signaling environments within motile cilia and flagella.

It is well-established that calcium signaling itself plays a central role in ciliary biology. Intraciliary calcium modulates dynein motor activity and ciliary beating, while calmodulin-dependent pathways contribute to centrosomal organization and basal body function [45–47]. In this context, the calcium-triggered liberation of a calmodulin-interacting globin domain with preferential centrosomal localization raises the possibility that ADGB participates in integrating calcium signaling with ciliary regulatory pathways. Although the precise biochemical activity of the liberated globin domain remains unresolved, the combined findings support a model in which ADGB functions as a dynamically regulated signaling protein whose proteolytic processing modifies its intracellular localization and interaction landscape.

A notable aspect of the current study is that the mechanistic and localization analyses were primarily performed in HEK293T cells using ectopic expression systems. These experimental approaches were intentionally chosen to enable controlled dissection of ADGB proteolysis, systematic analysis of truncation constructs, and biochemical characterization of this unusually large multidomain protein. Given the generally low endogenous expression levels of ADGB outside highly specialized ciliated and flagellated cell types [25], such controlled cellular systems currently provide one of the few experimentally tractable strategies to interrogate ADGB processing and domain-specific properties. Within this setting, the observed calcium-dependent proteolysis proved remarkably robust and reproducible across multiple experimental settings, including transient overexpression, CRISPR-based knock-in models, CRISPR activation of endogenous ADGB, and cell-free *in vitro* proteolysis assays. Simultaneously, it will be important for future studies to determine to what extent calcium-dependent ADGB processing and globin domain redistribution occur under endogenous physiological conditions in differentiated ciliated cell systems and *in vivo*. In particular, endogenous tagging approaches and advanced imaging in motile ciliated epithelia will be valuable to establish the temporal dynamics, cellular specificity, and physiological significance of ADGB proteolysis within its native biological context.

In conclusion, the present study identifies calcium-dependent proteolytic processing as a central and previously unrecognized feature of ADGB biology. We demonstrate that ADGB cleavage is mediated through a CAPNS1-associated calcium-dependent mechanism that generates stable cleavage products and liberates a globin-containing module with altered biochemical and cellular properties. Proteolytic processing enhances calmodulin interaction and enables preferential localization of the globin domain to centrosomal structures, thereby linking ADGB cleavage to pathways involved in calcium signaling and ciliogenesis. Together with recent structural and biochemical studies demonstrating atypical nitric oxide-related properties of the ADGB globin domain, our findings support a model in which ADGB acts as a dynamically regulated signaling globin within ciliated and flagellated cells. These observations provide a foundation for future studies aimed at dissecting the role of ADGB in calcium-responsive signaling and motile cilia function.

## Materials and Methods

### Cloning of recombinant human *ADGB* gene and the construction of ADGB mutants

Full length human ADGB and fragments of the gene were amplified from pLenti6-nGFP-ADGB for the generation of ADGB full length, deletion mutants or truncates. All plasmids were cloned into pFLAG-CMV^TM^-6a expression vector (Sigma-Aldrich®), unless otherwise indicated. An overview of mutant constructs can be found in **Supp. figure 4**. Detailed descriptions of the cloning protocol can be found in the supplementary methods. All ligation was carried out in temperature-cycle fashion in thermal-cycler programmed for 8 hours to cycle between 30 seconds at 10 °C and 30 seconds at 30 °C with 12 cycles per hour (modified from [48]), following heat-inactivation at 65 °C for 20 minutes. Ligated plasmids were transformed into DH5-α (*E. coli*) chemically competent cells.

### HEK293T cells culture, transfection and RNAi

HEK293T cells were cultured and maintained in DMEM (ThermoFisher) medium supplemented with 10% fetal bovine serum (Chemie Brunschwig) and penicillin-streptomycin mixed solution (penicillin: 100 units mL^-1^, streptomycin: 100 μg mL^-1^, ThermoFisher), DMEM/FBS/PS for simplification, and incubated at 37°C in a humidified incubator with 5% CO_2_. Unless otherwise stated, all transient expressions in this study were carried out with the calcium-phosphate precipitation method of transfection. Prior to transfection, HEK293T cells were seeded on 6-well plate(s) at a density of 5.2 x 10^4^ cells per cm^2^ of the dish surface area and were allowed to grow at 37°C/ 5% CO_2_ for 24 hours. At 60% - 80% of cell confluency, medium was aspirated and replaced with DMEM/FBS/PS containing 25 µM chloroquine (Sigma-Aldrich). In each well, 10% (v/v) of transfection mixture was added to the medium, consisting of 50% (v/v) 1.2 µg plasmid DNA/ 250 mM CaCl_2_ and 50% (v/v) HBS buffer (50 mM HEPES, 280 mM NaCl, 10 mM KCl, 1.5 mM Na_2_HPO_4_, 12 mM glucose). Transfected cells were incubated at 37°C/ 5% CO_2_ for 24 hours before the medium was replaced with fresh DMEM/FBS. Cells were harvested 48 hours post-transfection. For gene knock-down experiments, the same protocol was used with the addition of 100 pmol of esiEGFP/ esiCAPN1/ esiCAPN2/ esiCAPNS1 (Sigma-Aldrich) in a 6-well format, with or without the addition of plasmid DNA.

### Experimental hypoxia and oxidative stress

Oxygen deprivation on cell culture was induced by incubating the cells in a hypoxia chamber (Baker Ruskinn, InvivO_2_ 400) with a controlled environment of 0.2% O_2_, 5.0% CO_2_, 60% air humidity, at 37°C. For transient overexpression experiments, transfected HEK293T cells were initially incubated under normoxic conditions at 37°C/ 5% CO_2_ for 24 hours post-transfection to allow for cell recovery and protein expression, after which the cells were deprived of oxygen by incubation in the hypoxia incubator chamber. Cells were maintained in hypoxic environment for 24 hours prior to harvesting. For oxidative stress induction, HEK293T cells that were pre-incubated 24 hours in the hypoxia chamber were transferred and incubated under normoxic conditions for 1 hour prior to harvesting, in order to stimulate the production of reactive oxygen species. Validation of hypoxia and reoxygenation was done through detection and subsequent lack of detection of HIF-1α via immunoblotting.

### Treatment of calcium and pharmacological inhibitors

Calcium stimulated ADGB proteolysis was carried out by treating the transfected HEK293T cells with 50 mM CaCl_2_ alongside with 25 μM calcium ionophore A23187 (Sigma-Aldrich) for 1 hour prior to cell lysis. Pharmacological intervention of calcium-mediated ADGB proteolysis was carried out by treating the cells with inhibitors W-7 (Brunschwig, at 25 μM), MG132 (10 μM), calpeptin (Sigma-Aldrich, 25-100 μM), or PD150606 (Sigma-Aldrich, 5 – 25 µM) at the indicated concentrations for 23 hours prior to CaCl_2_/ calcium ionophore treatment.

### dCas9-VPR mediated expression of endogenous ADGB in HEK293T cells

The transcriptionally suppressed or silent *ADGB* gene in HEK293T cells was activated by employing the nuclease-null-Cas9 (of *Streptococcus pyogene*) with tandem fusion of VP64-p65-Rta tripartite activator (dCas9-VPR) [49] as described before [24] along with *ADGB* promoter-targeting guide RNAs (gRNAs). *ADGB* promoter-targeting gRNAs (gRNA1, 5’-GCTCTCCGGGCGCTGGACGC -3’; gRNA2, 5’- TTGCGTCCCTCTGCAGCCAC -3’) were cloned into and expressed from pSPgRNA plasmid (Addgene #47108). For dCas9-VPR activation of *ADGB* gene expression, HEK293T cells were transfected using Roti^®^-Fect (ROTH) method of plasmid DNA delivery, according to manufacturer’s instruction. Cells were transfected in 6-well plates, seeded with 5.2 x 10^4^ cells per cm^2^ surface area, 24 hours prior to transfection. In each well, 1125 ng of dCas9-VPR was delivered together with 375 ng of each gRNA1 and gRNA2. Cells were grown for 24 hours before the transfection medium was replaced with fresh DMEM/FBS/PS and allowed to grow for another 24 hours before CaCl_2_/ calcium ionophore treatment, as described above.

### RNA extraction and quantitative PCR

Total cellular RNA was extracted as previously described [50]. Total RNA (2 μg) was reverse transcribed (RT) using the Prime Script RT reagent kit (Takara Bio USA) and cDNA levels were estimated by quantitative polymerase chain reaction (qPCR) using the primers listed in **Supp. Table 2** and a KAPA SYBR® FAST qPCR reagent kit (Sigma-Aldrich) in a CFX96 C1000 Thermal Cycler (BioRad). Transcript levels were calculated as described before [51] and displayed as relative expression levels.

### Cell lysis and immunoblotting

For protein extraction, cells were gently washed with ice-cold phosphate-buffer-saline (PBS) buffer and a non-denaturing lysis buffer (10 mM Tris, pH 8.0, 1 mM EDTA, 400 mM NaCl, 0.1% NP-40, 2 μg mL^-1^ leupeptin, 2 μg mL^-1^ pepstatin, 2 μg mL^-1^ aprotinin, 1 mM PMSF) was added, cells were scraped, transferred to 1.5 mL microcentrifuge tubes and lysed on ice for 10 minutes. The lysate was centrifuged at 14,000 RPM, 4°C for 10 minutes. The concentration of soluble protein fraction was measured using Bradford (reagent from Brunschwig) protein assay as described previously [52].

Soluble proteins were separated with sample loading buffer (50 mM Tris-HCl, pH 6.8, 2% (w/v) SDS, 10% (v/v) glycerol, 100 mM DTT, 0.004% (w/v) bromophenol blue) in denaturing resolving gels (83 mm x 50 mm x 1.5 mm) consisting of linear 7.5%, 10.0% or 12.5% (w/v) polyacrylamide/N,N’-methylene-bis-acrylamide (AppliChem, 37.5:1), 375 mM Tris-HCl, pH 8.8, and 0.1% (w/v) SDS. The separated proteins were subsequently transferred onto nitrocellulose membrane (Amersham™ Protran® Western blotting membranes) in Towbin buffer (25 mM Tris-HCl, 192 mM glycine, 20% (v/v) ethanol) at 100V for 2 hours. After transfer, the membrane was blocked for 1 hour with 5% (w/v) skim milk in TBST buffer (25 mM Tris-HCl, pH 7.4, 150 mM NaCl, 0.1% Tween-20) at room temperature, followed by overnight incubation at 4°C with primary antibodies in 1% skim milk/TBST. The membrane was then washed three times with TBST buffer (8 minutes each), followed by incubation with secondary antibody (Sigma-Aldrich), for 1 hour at room temperature and washed for three times with TBST (8 minutes each). The membrane was dried and fluorescent signals were visualized Li-COR Odyssey^®^ infrared imaging system as described previously [53]. A list of antibodies is found in the supplementary methods.

### Purification of FLAG-GST-tagged ADGB

FLAG-GST-ADGB was produced in HEK293T cells for purification. HEK293T cells were cultured on 15-cm dishes and transfected using calcium phosphate precipitation method as described above. In each 15-cm dish, 20 µg of pFLAG-GST-ADGB plasmid was transfected and the cells were harvested 48 hours post-transfection for purification. Culture medium was aspirated and the cells were gentle washed with ice-cold PBS to remove excess medium. Triton-X lysis buffer (20 mM HEPES-NaOH, pH 7.5, 3 mM MgCl_2_, 100 mM NaCl, 1 mM EDTA, 1% Triton-X, 1 mM DTT, 1 mM PMSF) was then added to the dish and the cells were scraped and transferred into micro-centrifuge tubes, and incubated on ice for 10 minutes to allow cell lysis. Cell lysate was centrifuged at 14,000 RPM, 4°C for 10 minutes. The supernatant was filtered through 0.45-μm filter and incubated with Glutathione Sepharose® 4B (GSH4B) (GE Healthcare) (pre-washed with Triton-X lysis buffer) for 90 minutes at 4°C with rotation. After the incubation, GSH4B resin was centrifuged at 2000 RPM, 4°C for 2 minutes and the supernatant was discarded. The resin was washed with Washing buffer (20 mM Tris, pH8.0, 200 mM NaCl, 1 mM EDTA, 1 mM DTT, 2 mM ATP, 10 mM MgSO_4_) for 4 times and once with TED buffer (20 mM Tris, 1 mM EDTA, 1 mM DTT). Bound FLAG-GST-ADGB was released from the resin by incubation with Elution buffer (50 mM Tris, pH7.5, 1 mM EDTA, 100mM NaCl, 0.1% Triton-X, 80 mM Glutathione) at 4°C for 2 hours, with rotation. Following centrifugation at 2000 RPM, 4°C for 2 minutes, the eluted protein was recovered in the supernatant and dialysed in TED buffer using dialysis kit (GE Healthcare) at 4°C for 20 hours. Purified FLAG-GST-ADGB was collected, and the protein concentration was measured by Bradford protein assay. Purified protein was snap-frozen in liquid nitrogen and stored at -80°C until use.

### Extraction of cytoplasmic and nuclear fractions

Cytoplasmic and nuclear fractions were isolated from HEK293T cells with a protocol adapted from [54]. HEK293T cells were cultured on 15-cm dishes until confluency. After the removal of medium and gentle washing with ice-cold PBS, cells were lysed in Lysis buffer A (10 mM HEPES. pH 7.4, 10 mM KCl, 1.5 mM MgCl_2_, 0.5% NP-40, 1 mM DTT, 1 mM PMSF). Cells were scraped in Lysis buffer A, transferred to a centrifuge tube, and incubated on ice for 20 minutes. The cell lysate was centrifuged at 14,000 RPM, 4°C for 10 minutes and the supernatant was recovered as the cytoplasmic fraction. The pellet was then resuspended in Lysis buffer B (20 mM HEPES, pH 7.9, 400 mM NaCl, 1.5 mM MgCl_2_, 0.2 mM EDTA, 15% glycerol, 1 mM DTT, 1 mM PMSF) and incubated at 4°C for 20 minutes with rotation. The resuspended pellet was sonicated and centrifuged at 14,000 RPM, 4°C for 10 minutes. The supernatant recovered as the nuclear fraction. To assess the purity of each fraction, 50 μg were resolved on a 7.5% denaturing polyacrylamide gel and immunoblotted against tubulin (cytoplasmic content) and lamin B (nuclear content).

### *In vitro* cleavage of ADGB

For *in vitro* cleavage of ADGB in cells lysate, HEK293T cells transiently expressing FLAG-ADGB-myc or EGFP-ADGB were lysed in non-denaturing lysis buffer with or without the chelating agent EDTA. 100 μg of total protein lysate was incubated at 37°C for 0, 1, 3, 6, or 24 hours. The reaction was stopped by adding 5X sample loading buffer and heating to 95°C for 5 minutes.

For *in vitro* cleavage of purified ADGB, purified FLAG-GST-ADGB (∼1.2 μg) was reconstituted in TED buffer and incubated at 37°C for 22 hours. To examine the effect of calcium-ions on ADGB cleavage, purified FLAG-GST-ADGB was incubated with or without 10 mM CaCl_2_. Subsequently, FLAG-GST-ADGB was also incubated with 35 μg of cytoplasmic or nuclear extract, with and without the addition of 10 mM CaCl_2_. Similarly, all reactions were stopped by adding 5X sample loading buffer and heating to 95°C for 5 minutes. The presence of cleaved ADGB was determined by immunoblotting against ADGB.

### Immunofluorescence and microscopy

To investigate the localization of full-length ADGB and its truncated mutants, plasmid DNA coding for the respective GFP-tagged proteins were overexpressed in HEK293T cells cultured on glass cover slip, using the calcium phosphate precipitation method as described above. Cells were fixed with 3.7% paraformaldehyde/PBS for 10 minutes at RT 48 hours post-transfection. Fixing solution was completely removed followed by washing with 3 mL of PBS solution. Then, cells were permeabilized with 1 mL of 10% Triton-X/PBS, on ice for 15 minutes and blocked with 1 mL of 10% FBS/PBS at room temperature for 1 hour. Subsequently, the 10% FBS/PBS was removed and the cells were washed with 3 mL of PBS solution for 1 hour. The cells were then incubated overnight with mouse anti-acetylated tubulin (Santa Cruz) or anti-PCM-1 (Proteintech) diluted at 1/200 in 10% FBS/PBS. After overnight incubation, cells were washed in ice-cold PBS for 1 hour, followed by incubation with secondary Alexa Fluor goat anti-mouse or anti-rabbit IgG (Invitrogen) diluted 1/1000 in PBS. Cells were again washed in PBS for 1 hour and carefully mounted on a microscope slide with a Fluoromount-G^TM^, with DAPI (SouthernBiotech). The fluorescent signals were detected and imaged on a Nikon Eclipse fluorescent microscope (Nikon Corporation) or Leica SP5 confocal microscope (Leica Microsystem).

### Mammalian one-hybrid system

In order to measure the potential proteolytic activity of ubiquitous calpains, notably CAPN1 and CAPN2, the mammalian one-hybrid system was employed. In this system, a short peptide containing the calpain proteolytic cleavage site was cloned between a Gal4 DNA-binding domain and VP16 transcriptional activator domain. Employing luciferase-based assays, Gal4-VP16 fusion protein binds on concatemerized Gal4 responsive element-driven E1b promoter-dependent *Luciferase* gene and increases the luciferase signal. In contrast, the separation of Gal4 and VP16 domains via proteolytic cleavage by calpains results in a diminished luciferase signal. Here, the human spectrin (hSpec) cleavage site was cloned between Gal4 and VP16 (see supplementary methods).

HEK293T cells were seeded with 5.2 x 10^4^ cells per cm^2^ surface area 24 hours prior to transfection using calcium phosphate precipitation method of plasmid DNA delivery, as described above. After 24 hours of incubation, medium was aspirated and replaced with DMEM/FBS without antibiotics. In this experiment, cells were transfected with 250 ng of pGRE5xE1b, 250 ng of pM3-Gal4-hSpec-VP16, along with 5 ng of pRL-SV40 *Renilla* luciferase control plasmid. Cells were treated with 5 μM, 10 μM, and 25 μM of PD150606, 6 hours after transfection with subsequent incubation of 16 hours. Cells were then lysed with Passive Lysis Buffer (Promega, Wisconsin, USA) and the luciferase activities of soluble lysate were measured using the Dual Luciferase Reporter Assay System as previously reported[53].

### Generation of *ADGB* knock-in cell models

CRISPR-based *ADGB* knock-in was performed by targeting and inserting the *ADGB* coding sequence into AAVS1 site in human chromosome 19 [55]. Full length *ADGB* coding sequence was cloned into pAAVS1-P-CAG-DEST vector, which is a generous gift from Knut Woltjen (Addgene plasmid # 80490), through the Gateway recombination according to the manufacturer’s protocol, to generate pAAVS1-P-CAG-ADGB-DEST. Cloned *ADGB* coding sequence is situated downstream of a CMV promoter and Puromycin selection gene, flanked by AAVS-left and -right homology arms. A partnering plasmid pXAT2, which is a generous gift from Knut Woltien (Addgene plasmid # 80490), was co-transfected as the template for AAVS1 sgRNA and Cas9 protein. One day prior to transfection, HEK293T cells, MCF-7 cells, and HeLa cells were seeded on 6-well plates at 5.2 x 10^4^ cells per cm^2^ surface area. Cells were then transfected with 1.2 μg pAAVS1-P-CAG-ADGB-DEST and 1.2 μg pXAT2 using calcium phosphate method of transfection for HEK293T cells (as described above) and ROTI^®^Fect for MCF-7 cells and HeLa cells (according to manufacturer’s protocol). Upon confluency, cells were subjected to puromycin selection at 2.0 μg/mL for HEK293T and HeLa cells and 1.0 μg/mL for MCF-7 cells. Cells were selected for at least two weeks before analysis and continuously maintained under puromycin. *ADGB* knock-in was verified via PCR of genomic DNA using primer pair flanking a region within the *ADGB* gene (5’-TGGAGGAGTGTCTTCACCAG-3’), and the AAVS1-right arm (5’-GTCAGAGCAGCT CAGGTTCT-3’). Immunoblotting and immunostaining against ADGB were also performed for verification.

### Immunoprecipitation

Immunoprecipitation was carried out in lysates of HEK293T cells with prior transient overexpression of bait and prey proteins. Cell lysates were prepared using NP-40 lysis buffer (20 mM HEPES/NaOH (pH7.5), 3 mM MgCl_2_, 100 mM NaCl, 1 mM DTT, 1 mM EGTA, 0.5% NP-40, 2 μg mL^-1^ leupeptin, 2 μg mL^-1^ pepstatin, 2 μg mL^-1^ aprotinin, 1 mM PMSF, 1 mM DTT). Cell lysates were subjected to 14,000 RPM centrifugation to recover the supernatant, followed by incubation with 2 µg of primary antibody: anti-GFP (Proteintech), anti-ADGB (Sigma-Aldrich), or anti-rabbit IgG (GE healthcare), at 4°C for 2 hours. The lysates with antibodies were then centrifuged at 14,000 RPM before transferring to 30 µL (bed volume) of Protein G Sepharose® 4 Fast Flow (Cytiva) resin and incubated with rotation at 4°C for 2 hours. The resin was then washed 5 times with wash buffer (20 mM HEPES/NaOH (pH7.5), 3 mM MgCl_2_, 150 mM NaCl, 1 mM DTT, 1 mM EGTA, 0.5% NP-40, 2 μg mL^-1^ leupeptin, 2 μg mL^-1^pepstatin, 2 μg mL^-1^ aprotinin, 1 mM PMSF, 1 mM DTT). To elute the immunoprecipitated proteins, the resin was boiled in equal volume of 2X sample loading buffer (50 mM Tris-HCl, pH 6.8, 2% (w/v) SDS, 10% (v/v) glycerol, 100 mM DTT, 0.004% (w/v) bromophenol blue). The immunocomplexes were analyzed by immunoblotting.

## Data availability

All data are given in the manuscript.

## Supporting information

## Acknowledgments

We thank Antonia Herwig for assistance with figure preparation and all Hoogewijs lab members for critical reading of the manuscript.

## Funding and additional information

This work was supported by the Swiss National Science Foundation to D.H. (project nr. 310030_207460).

## Author contributions

T.W.K. and D.H. conceptualization; T.W.K., C.O. and A.C. investigation; T.W.K. visualization; T.W.K., C.O. and D.H. writing–original draft; D.H. project administration; D.H. resources; D.H. funding acquisition; D.H. supervision.

