## Supplemental methods and figures for "Calcium-dependent proteolysis of the chimeric androglobin reveals altered localization of its isolated globin domain"

**Running title:** Androglobin undergoes calcium-dependent proteolysis

### Supplementary Methods
### Supplementary Tables
### Supplementary Figures and legends

### Supplementary Methods

#### Cloning of recombinant human ADGB gene and the construction of ADGB mutants

A doubly-tagged recombinant ADGB clone was constructed by the amplification of full-length *ADGB* by PCR with a pair of primers (forward, 5'-AAAAAGCAGGGTCGACCATGGCCTC CAAACAAACCAAAAAG-3', *RE* site underlined; and reverse, 5'-GAAGGCACAG GGTACCTTACAGGTCCTCTTCTGAGATCAGCTTCTGCTCAAAGCTGGGTCTAGATG CCTTTTTCTTTCC-3', *RE* site, underlined, with an additional c-myc sequence included). The amplicon was then digested and ligated into linearized pFLAG-CMV$^{TM}$-6a vector, to generate pFLAG-ADGB-myc. Full-length ADGB used in this study lacks 6 amino acids (res. 26 - 31) at the N-terminal compared to the RefSeq version NM_024694.4. This is highly likely to be arisen from the splicing event as the 6 amino acids are located at the exon-exon splice junction. For consistency, the numbering of ADGB residues will be based on the full-length ADGB sequence as deposited in UCSC Genome Browser. Halo-tag ADGB was generated by amplifying ADGB coding sequence using primer pairs (forward, CAAAAAAGCAGCGATCGCCATGGCCTCC AAACAAACCAAAAAG, *RE* site underlined; reverse, TCAACCACTTGTTTAAACAAAG CTGGGTCTAGATTACTTTTTCTTTCC, *RE* site underlined), and cloned into pFN21A HaloTag® CMV Flexi® Vector (Promega), according to manufacturer's protocol.

ADGB deletion mutants lacking different domains, the IQ domain (ΔIQ), parts of the uncharacterized 350-residue domains (ΔPro.350res. and ΔDis.350res) and C-terminal domains (ΔPro.C-ter and ΔDis.C-ter), were constructed by ligating amplicons from two parts of the *ADGB* gene with custom designed complementary overhangs. The N-terminal fragments were amplified by PCR using primer pairs (forward, 5'-AAAAAGCAGGGTCGACCATGGCCTCC AAACAAACCAAAAAG-3', *RE* site underlined; and reverse 5' – AATTGCTGCT GGTCTCCTGGCAGCAGAATACTCTTTATCACTTGTGGGCATAC – 3' for ΔIQ, 5' – CTG ATGAGCAGGTCTCAATCCTGCAGAAGAACAATCTGATAGGGACTGC – 3' for ΔPro.350res., 5' – AATAT ATATAGGTCTCATGTGTACACAGAAATCTTCAAAATCTA ACC – 3' for ΔDis.350res., and 5'- ACGGCATGGTGGTCTCGAGCAATCTTAGTTTCTTC ATCTTG -3' for ΔPro.C-ter, *RE* site underlined). Similarly, the C-terminal fragments were amplified by PCR using primer pairs (forward, 5'- CAGATTGTTCGGTCTCAATACAGACC TCTCATATGGTCGTATATGC-3' for ΔProtease, 5' – GCTATCAAGATGGGTCTCCCAT AATTGCTTTTGCAGATTATACTGTGAC – 3' for ΔGlobin , 5' – TTAGATTGCTGGTCTC AGCCAGAATACCAGACACAAAAGAAAATATCAGTG – 3' for ΔIQ, 5' – ATGAACAAA

CGGTCTCTGGATTGGGTGATGCTCATCAGAGTGATG – 3' for ΔPro.350res., 5' – CAGTAGGACAGGTCTCACACATCTGCAGCATGGTGTC – 3' for ΔDis.350res., and 5'- TATGGTGAAAGGTCTCATGCTTTAATTAACTTAGGAAGCCCAGACTCC -3' for ΔPro.C-ter, *RE* site underlined; and reverse 5'-GAAGGCACAGGGTACCTTACAGGTCCTCTTCTGAGATCAGCTTCTGCTCAAAGCTGGGTCTAGATGCCTTTTTCTTTCC-3', *RE* site underlined, with an additional c-myc sequence included). For ΔDis.C-ter, single amplicon was amplified by PCR using the primer pairs (5'-AAAAAGCAGGGTCGACCATGGCCTCCAAACAAACCAAAAAG-3' *RE* site underlined; and reverse, 5'- AAGATTTTGGGTACCTTACAGGTCCTCTTCTGAGATCAGCTTCTGCTCGGCGGCCTCCTCGTGTCTTTCACCATAAGC -3', *RE* site underlined, with an additional c-myc sequence included). These amplicons were then digested with their respective restriction enzymes as designed on the primers, and ligated (2- or 3-way ligation) into linearized pFLAG-CMV$^{TM}$-6a vector digested with *Xho*I and *RE*, to generate pFLAG-ADGBΔProtease-myc (res. 57 – 409 removed), pFLAG-ADGBΔGlobin-myc (res. 763 – 985 removed), pFLAG-ADGBΔIQ-myc (res. 905 – 927 removed), pFLAG-ADGBΔPro.350res-myc (res. 410 – 588 removed), pFLAG-ADGBΔDis.350res-myc (res. 641 – 762 removed), pFLAG-ADGBΔPro.C-ter-myc (res. 988 – 1295 removed), and pFLAG-ADGBΔDis.C-ter-myc (res. 1296 – 1667 removed). All deletion mutants contain a FLAG-tag and a c-myc-tag at the N-terminus and the C-terminus, respectively. Generation of ADGB-ΔIQ and ADGB-Δcalpain constructs were described before (Keppner et al. 2022)

ADGB truncated mutants consisting of only the N-terminal domain (res. 1 – 57) and the C-terminal domain (res. 969 – 1667) were cloned. These truncates were designed to carry a FLAG-GFP tag at the N-terminal and GFP tag at the C-terminal with glycine-serine linkers (GGSGGGGSGG) between the tags and the ADGB domains. The N-terminal GFP tag was amplified by PCR using a primer pair (forward, 5'- GCTAGTTAAGTCGACCATGGTGAGCAAGGGCGAGGAG-3', *RE* site underlined; reverse, 5'- ATATAGGTCTCA**TCCTCCACTACCACCACCACCACTACCTCC**CTTGTACAGCTCGTCCATGCCGAG, *RE* site underlined, glycine-serine linker in bold). Similarly, the C-terminal GFP tag was amplified using a primer pair (forward, 5'- GACTCGGTCTCG**GGAGGTAGTGGTGGTGGTGGTAGTGGAGGA**ATGGTGAGCAAGGGCGAGGAG -3', *RE* site underlined, glycine-serine linker in bold; reversed, 5'- TGTACAAACTGGTACCTTACTTGTACAGCTCGTCCATGCCGAG -3', *RE* site underlined). Truncates of ADGB domains with complementary overhangs to the above N- and C-terminal tags were amplified using primer pairs, for N-terminal domain (forward, 5'- CACAAGGGTCTCCAGGAATGGCCTCCAAACAAACC-3'; reverse, 5'- GCA TCCGGTCTCTCTCCATTTATGTCAGCTTCACTCC -3', *RE* site underlined), and the C-terminal domain (forward, 5'- GACTAAGGTCTCAAGGAAAGTGCAAGTCTTTGGAATC -3'; reverse, 5'- GAGATCGGTCTCTCTCCAAAGCTGGGTCTAGATGCC -3', *RE* site underlined). These amplicons were then digested with their respective restriction enzymes as designed on the primers, and ligated (4-way ligation) into linearized pFLAG-CMV™-6a vector, to generate pFLAG-GFP-Nter-GFP (res. 1 – 57, without res. 26 – 31) and pFLAG-GFP-Cter-GFP (res. 969 – 1667).

Single GFP-tagged full length ADGB, globin domain (ADGB_GD), and globin domain with C-terminal segment (ADGB_GD-Cter) were cloned by using the above mentioned N-terminus GFP-tagged to ligate with amplicons amplified by PCR using the primer pairs for full length ADGB (forward, 5'- CACAAGGGTCTCCAGGAATGGCCTCCAAACAAACC-3, *RE* site underlined; reverse, reverse, 5'-GAAGGCACAGGGTACCTTACAGGTCCTCTTCTGAGA TCAGCTTCTGCTCAAAGCTGGGTCTAGATGCCTTTTCTTTCC-3', *RE* site underlined), ADGB_GD (forward, CAGTAGGGTCTCCAGGACACATCTGCAGCATGGTGTC, *RE* site underlined; reverse, ACCACCGGTACCTTAGCTTTTAAACATTAGTCTTAAGAGAGAA ACTGC, *RE* site underlined), and ADGB_GD-Cter (forward, CAGTAGGGTCTCCAGGAC ACATCTGCAGCATGGTGTC, *RE* site underlined; reverse, 5'-GAAGGCACAGGGT ACCTTACAGGTCCTCTTCTGAGATCAGCTTCTGCTCAAAGCTGGGTCTAGATGCCT TTTTCTTTCC-3', *RE* site underlined). Digested amplicons were ligated into linearized pFLAG-CMV™-6a vector to generate pFLAG-GFP-ADGB, pFLAG-GFP-ADGB_GD, and pFLAG-GFP-ADGB_GD-Cter. With the same construction to pFLAG-GFP-ADGB_GD, GFP-tagged ADGB globin domains with single mutation on the proximal histidine (H824G) or distal glutamine (Q792G), or both (H824G/Q792G) were cloned by amplifying and ligating the globin gene using primers designed to carry the mutated codon sequence.

FLAG-tagged ADGB N-terminal segment consisting of the res. 1 to 987 of the protein was cloned by ligating linearized pFLAG-CMV™-6a vector with digested amplicon amplified using primer pairs (forward, 5'- GCTAGTTAAGTCGACCATGGTGAGCAAGGGCGAGG AG-3', *RE* site underlined; reverse, ACCACCGGTACCTTAGCTTTTAAACATTAGTCTTA AGAGAGAAACTGC, *RE* site underlined), to generate pFLAG-Nter-ADGB_GD.

For protein purification, full-length ADGB with GST-tag at the N-terminus was constructed, with a glycine-serine linker (GGSGGGGSGG) between the GST-tag and the ADGB. The N-terminal GST-tag was amplified by PCR, with pDEST$^{TM}$20 vector (Invitrogen) as a template, using a primer pair (forward, 5'- CGTACGGTCGACCATGGCCCCTATACTA GGTTA -3', *RE* site underlined; reverse, 5'- ATATAGGTCTCA**TCCTCCACTACCACCACCACCAC TACCTCC**TTTTGGAGGATGGTCGCCAC -3', *RE* site underlined, glycine-serine linker in bold). Full-length ADGB was amplified with a primer pair (forward, 5'- AATAAGGGTCTC CAGGAATGGCCTCCAAACAAACCAAAAAG -3', *RE* site underlined, reverse, 5'- ACCA CTGGTACCTTAAAAGCTGGGTCTAGATGCCTTTTTCTTTCC -3', *RE* site underlined). These amplicons were then digested with their respective restriction enzymes as designed on the primers, and ligated (3-way ligation) into linearized pFLAG-CMV$^{TM}$-6a vector to generate pFLAG-GST-ADGB.

For the study of ADGB globin and IQ domain, ADGB mutant with R918Q/R923Q were synthesized in pENTR$^{TM}$4 vector, and the gene was sub-cloned into pcDNA3.1-EGFP Destination vector, according to manufacturer's protocol, to generate pcDNA3.1-EGFP-ADGB(IQ.RR) plasmid.

**Cloning of hSpec cleavage site**

The amplicon for hSpec fragment was generated from synthesized complementary oligo strands complexed in T4 DNA ligase buffer (ThermoScientific) (40 mM Tris-HCl, 10 mM $MgCl_2$, 10 mM DTT, 500 μM ATP) with T4 poly nucleotide kinase (ThermoScientific) at 37°C for 1 hour, followed by heating to 95°C for 5 minutes and slow cooling at the rate of -5°C $min^{-1}$ to 10°C, to produce oligo duplex (with *EcoR*I overhangs at both ends). The oligo duplex is then cloned into pM3-Gal4BD-VP16AD vector linearized with *EcoR*I digestion, generating pM3-Gal4-hSpec-VP16 (insert: *GSG*QQEVYGMMPRD*GSG*, hSpec underlined, glycine-serine linker *italized*).

**Antibodies**

The antibodies used in this study include anti-ADGB (1:500) (Sigma-Aldrich, HPA036340-100UL), anti-GFP (1:1000) (Proteintech, 50430-2-AP-150UL), anti-FLAG (1:1000) (Proteintech, 20543-1-AP (rabbit)/ 66008-3-IG-150UL (mouse)), anti-GST (1:500) (OriGene, TA150033), anti-tubulin (1:1000) (Santa Cruz, sc-23950), anti-laminB (1:1000) (abcam, ab16048), anti-HIF1α (1:1000) (BD Transduction Laboratories™, 610958), and anti-V5 (1:1000) (Proteintech, 14440-1-AP-150UL).

**Supplementary Table 1:** List of centrosomal and/or ciliary proteins co-immunoprecipitated with the isolated globin domain

| Protein | Enrichment in Glob-GFP (log2) | GO centrosome | GO_ciliary basal body | HPA location centrosome | HPA location basal body | Comments |
|---|---|---|---|---|---|---|
| **CCT8** | 6.08 | y | n | n | n | localisation to sperm connecting piece |
| **TCP1** | 4.68 | y | n | - | - | no data |
| **EMD** | 4.48 | y | n | n | n | |
| **RUVBL1** | 4.27 | y | y | n | y | |
| **CALM1** | 4.18 | y | n | n | n | |
| **BIRC6** | 3.98 | y | n | n | n | |
| **KIF11** | 3.90 | n | y | n | y | localisation to sperm midpiece |
| **ESPL1** | 3.76 | y | n | n | n | |
| **PCNA** | 3.69 | y | n | n | n | |
| **ALMS1** | 3.54 | y | y | y | y | |
| **BCAS2** | 3.39 | y | n | y | n | |
| **IQSEC1** | 3.20 | y | n | y | n | |
| **SLC16A1** | 3.14 | y | n | n | n | |
| **STIL** | 3.13 | y | y | y | y | |
| **SLC1A5** | 3.10 | y | y | y | y | localisation to centriolar satellite, uncertain |
| **RBM39** | 3.08 | y | n | y | n | localisation to centriolar satellite, uncertain |
| **RPAP3** | 3.01 | n | y | n | y | localisation to sperm acrosome |
| **SPAG5** | 2.92 | y | y | n | n | |
| **ERC1** | 2.90 | y | y | y | n | |
| **MMS19** | 2.84 | y | n | n | n | |
| **CEP85** | 2.61 | y | n | y | n | |
| **DNM2** | 2.52 | y | n | n | n | |
| **HAUS5** | 2.52 | y | n | - | - | no data |
| **USP9X** | 2.33 | y | n | n | n | |
| **NCAPD2** | 2.29 | y | y | y | y | localisation uncertain |
| **OBSL1** | 2.28 | y | n | y | n | |
| **PIK3R4** | 2.26 | n | y | n | y | localisation uncertain |
| **TUBGCP2** | 2.13 | y | y | y | y | |

| | | | | | | |
|---|---|---|---|---|---|---|
| **KRT18** | 2.11 | y | n | n | n | |
| **PSMB5** | 2.04 | y | n | y | n | localisation to sperm midpiece |
| **PSMA1** | 2.00 | y | n | y | n | localisation uncertain |
| **WDR62** | 1.95 | y | n | y | n | localisation to centriolar satellite |
| **CDK5RAP2** | 1.94 | y | y | y | y | localisation to sperm midpiece |
| **PKN2** | 1.86 | y | n | y | n | |
| **CCT5** | 1.80 | y | n | n | n | localisation to sperm midpiece |
| **CCT4** | 1.72 | y | n | n | n | |
| **HAUS6** | 1.65 | y | n | y | | localisation to centriolar satellite & sperm midpiece |
| **ACLY** | 1.60 | n | y | n | y | localisation to flagellar endpiece, uncertain |
| **DCTN1** | 1.56 | y | y | n | y | |
| **WDR11** | 1.48 | n | y | y | n | localisation to flagellar centriole & principal piece |
| **DYNC1H1** | 1.48 | y | n | n | n | localisation to flagellar centriole & midpiece |
| **DYNC1I2** | 1.42 | y | n | y | n | localisation to flagellar centriole & midpiece |
| **RRM1** | 1.39 | y | y | y | y | localisation to centriolar satellite, uncertain |
| **CTDP1** | 1.38 | y | n | n | n | |
| **YTHDF2** | 1.36 | y | n | n | n | cell-line dependent localisation to centriolar satellite |
| **LUZP1** | 1.26 | y | y | y | n | |
| **ATXN10** | 1.23 | n | y | n | n | |

| | | | | | | |
|---|---|---|---|---|---|---|
| **FLII** | 1.23 | y | n | y | n | localisation to centriolar satellite |
| **SKP1** | 1.18 | y | n | y | y | |
| **NEK9** | 1.17 | y | n | - | - | no data |
| **TUBGCP6** | 1.04 | y | n | - | - | no data |
| **KIF7** | 1.02 | n | y | n | n | |

### Supplementary Table 2. qPCR primers

| Target | Primer fwd | Primer rev |
|---|---|---|
| **ACTIN** | CTG GAA CGG TGA AGG TGA CA | AAG GGA CTT CCT GTA ACA ATG |
| **CAPN1** | AGA GTG GAA CAA CGT GGA CC | AAG GTG GCT GGG TAG TTT CG |
| **CAPN2** | AAGTAACGGAAGCCTACAGAAAC | ATCTTCATGCCGTCTGGTCAG |
| **CAPSN1** | CGT GAT GGA TAG CGA CAC CA | TCA TTC AGG TGG AAC CCT GC |

### Supplementary figures:

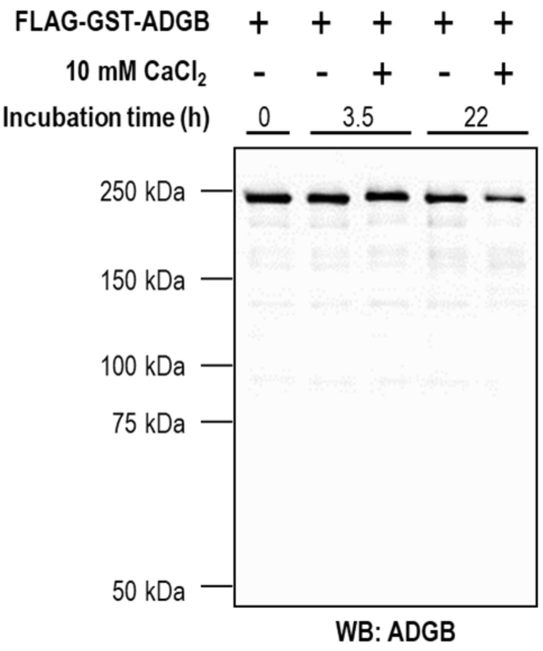


**Supplementary figure 1. Recombinant ADGB does not exhibit instantaneous cleavage in the presence of calcium.** *In vitro* autolysis assay of purified FLAG-GST-ADGB (~1.2 µg) with and without 10 mM $CaCl_2$, incubated at 37°C for 3.5 or 22 hours. Full-length ADGB remained stable and proteolytic products were not detected under all conditions.

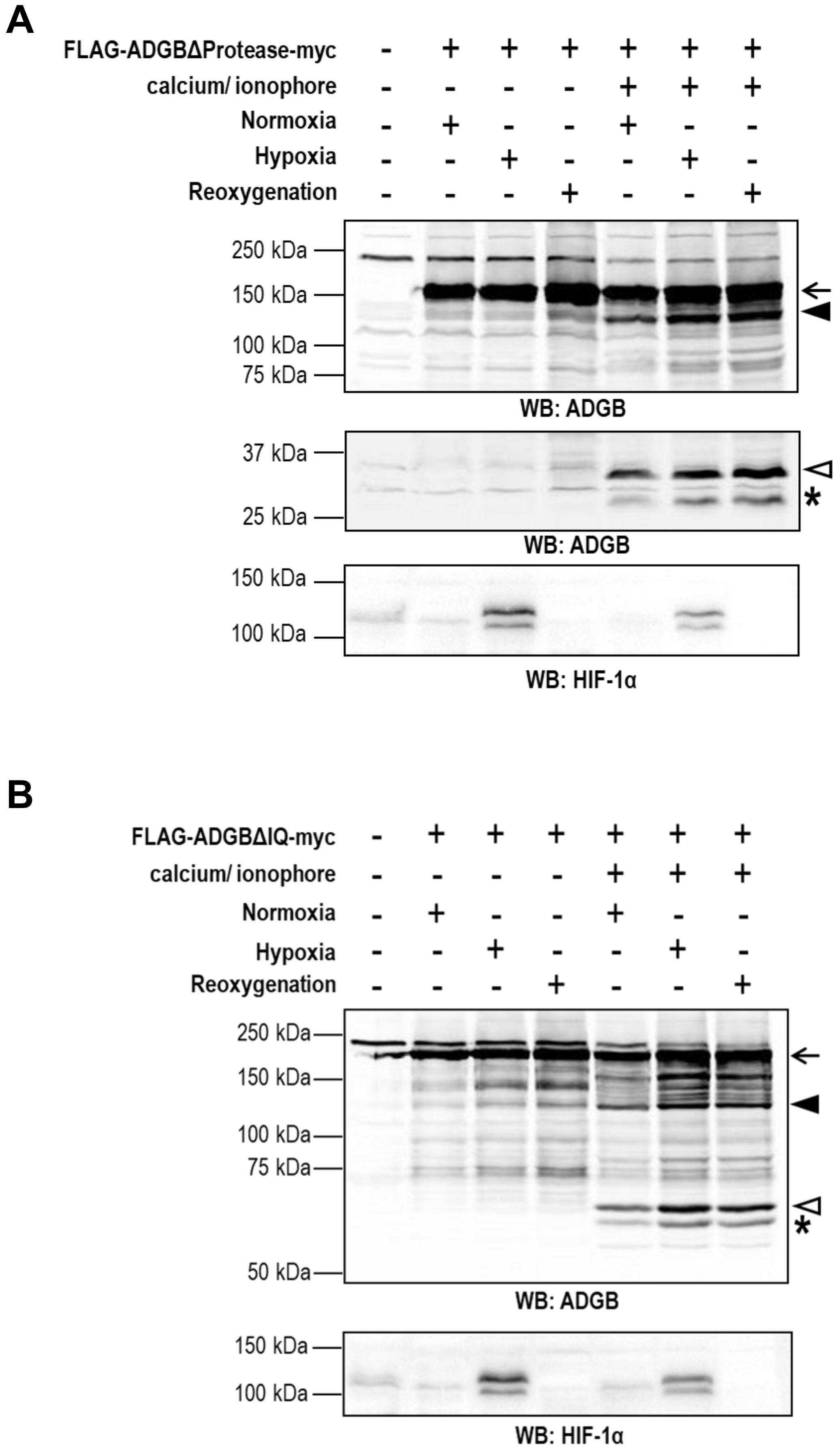
A
FLAG-ADGBΔProtease-myc
calcium/ ionophore
Normoxia
Hypoxia
Reoxygenation
250 kDa
150 kDa
100 kDa
75 kDa
WB: ADGB
37 kDa
25 kDa
WB: ADGB
150 kDa
100 kDa
WB: HIF-1α
B
FLAG-ADGBΔIQ-myc
calcium/ ionophore
Normoxia
Hypoxia
Reoxygenation
250 kDa
150 kDa
100 kDa
75 kDa
50 kDa
WB: ADGB
150 kDa
100 kDa
WB: HIF-1α

**Supplementary figure 2. ADGB protease- and IQ-motif deletion mutants exhibit no change in calcium-mediated cleavage under hypoxic conditions.** FLAG-ADGB-myc was transiently expressed in HEK293T cells in normoxia (atmospheric $O_2$ level), hypoxia (0.2% $O_2$), and hypoxia followed by reoxygenation (ROS stress), with subsequent analysis as described in Figure 4(A). The calcium-mediated proteolytic events remained unchanged under these conditions in both (A) protease domain deletion mutant and (B) IQ-motif deletion mutant.

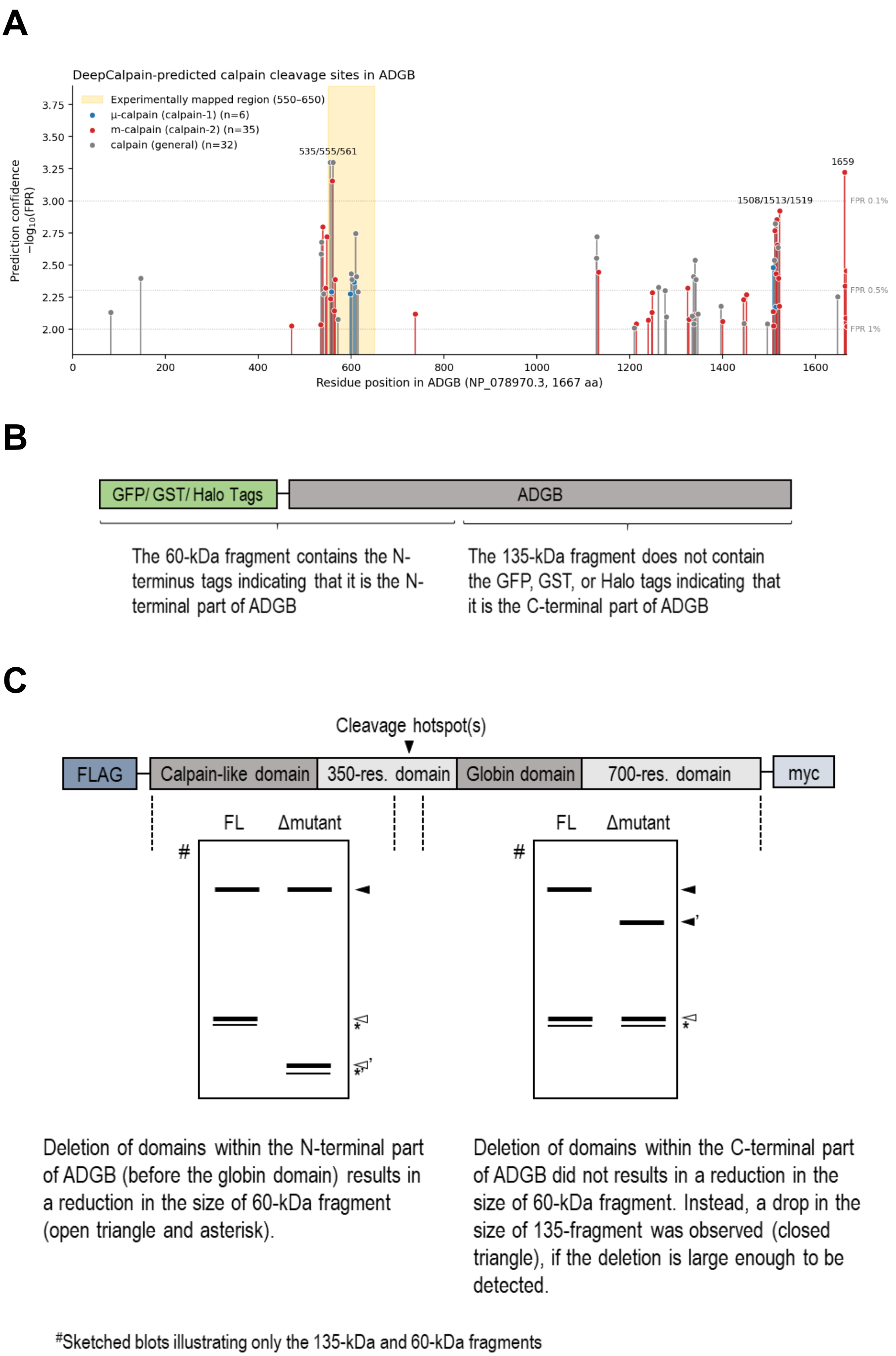

A
DeepCalpain-predicted calpain cleavage sites in ADGB
Experimentally mapped region (550–650)
μ-calpain (calpain-1) (n=6)
m-calpain (calpain-2) (n=35)
calpain (general) (n=32)
535/555/561
1508/1513/1519
1659
FPR 0.1%
FPR 0.5%
FPR 1%
Prediction confidence −log10(FPR)
Residue position in ADGB (NP_078970.3, 1667 aa)
B
GFP/ GST/ Halo Tags
ADGB
The 60-kDa fragment contains the N-terminus tags indicating that it is the N-terminal part of ADGB
The 135-kDa fragment does not contain the GFP, GST, or Halo tags indicating that it is the C-terminal part of ADGB
C
Cleavage hotspot(s)
FLAG
Calpain-like domain
350-res. domain
Globin domain
700-res. domain
myc
FL
Δmutant
#
Deletion of domains within the N-terminal part of ADGB (before the globin domain) results in a reduction in the size of 60-kDa fragment (open triangle and asterisk).
Deletion of domains within the C-terminal part of ADGB did not results in a reduction in the size of 60-kDa fragment. Instead, a drop in the size of 135-fragment was observed (closed triangle), if the deletion is large enough to be detected.
#Sketched blots illustrating only the 135-kDa and 60-kDa fragments

**Supplementary figure 3.** A. Predicted calpain cleavage sites displays a hotspot in amino-acids 550-650 of ADGB, corresponding to the experimentally mapped cleavage. B. Schematic representation of 60-kDa and 135-kDa fragments produced in calcium-mediated ADGB cleavage as observed in Figure 1A. C. Size-based prediction of calcium-dependent cleavage hotspot(s) in ADGB based on changes in the sizes of the 60-kDa and 135-kDa fragments.

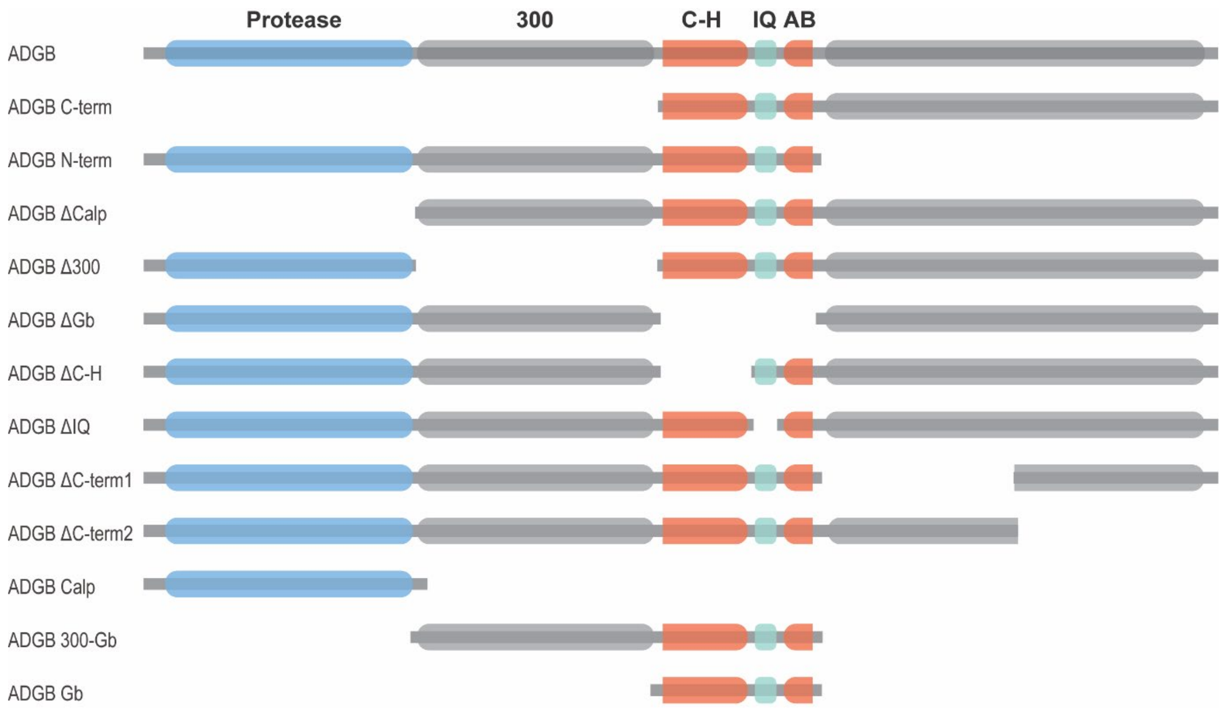


**Supplementary figure 4.** Schematic representation of ADGB mutants used in this study.

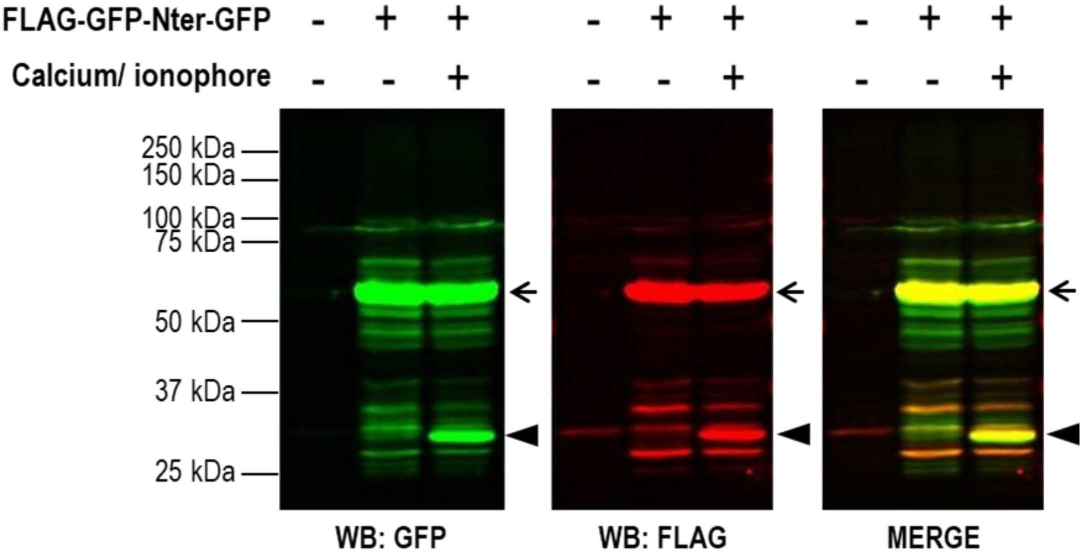


**Supplementary figure 5. Truncated mutant of the ADGB N-terminal domain might contain a calcium-dependent cleavage site.** The ADGB N-terminus segment flanked with FLAG-GFP at the N-terminus and GFP at the C-terminus was overexpressed in HEK293T cells with and without treatment of 50 mM $CaCl_2$/ 25 μM calcium ionophore. Immunoblotting analysis against GFP and FLAG shows calcium dependent proteolysis in this segment. Dual-mode imaging of GFP and FLAG was used to differentiate the N-terminal fragments (with FLAG tag) and the C-terminal fragment (without FLAG tag). The full-length protein and its cleaved product are indicated by arrow and closed triangle, respectively.

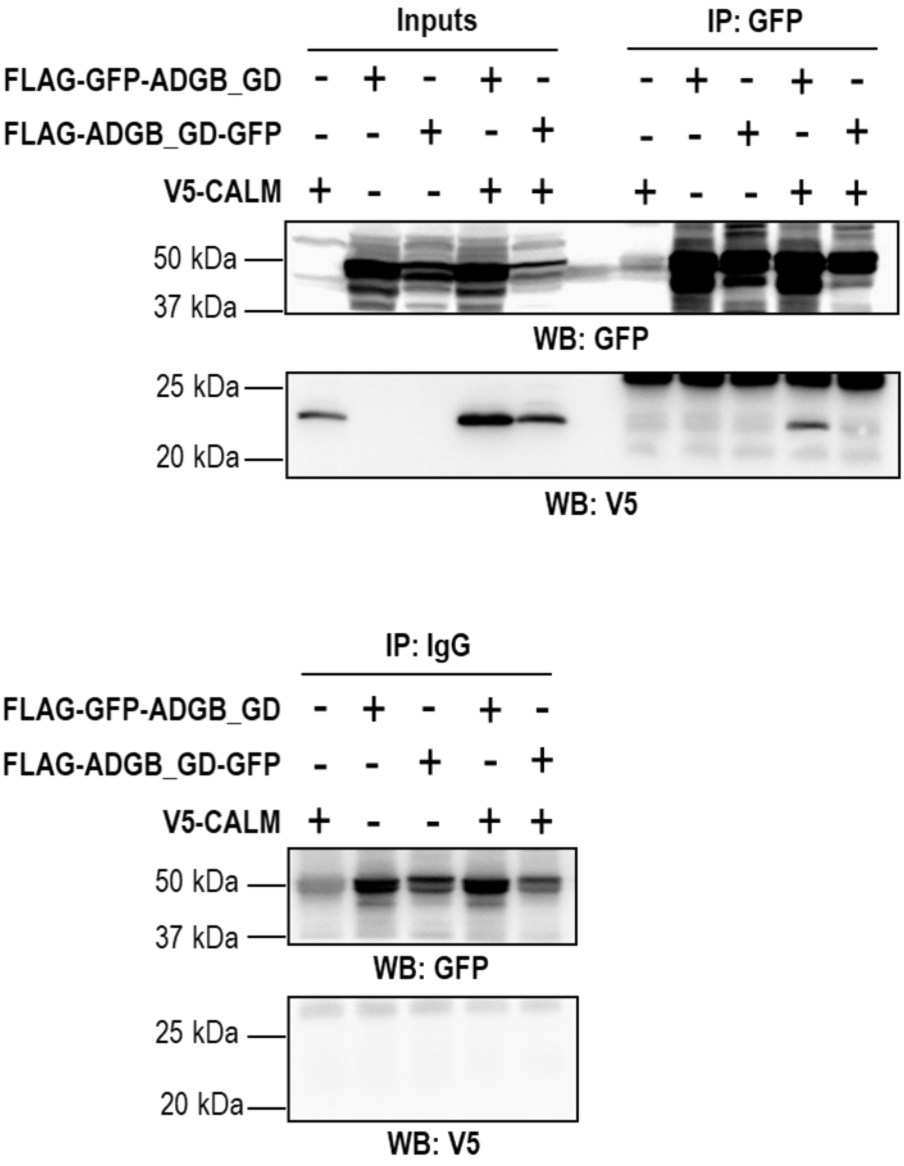


**Supplementary figure 6. ADGB globin domain interacts with calmodulin and the interaction is interfered when GFP is situated C-terminal to the globin domain.** Co-immunoprecipitation of FLAG-GFP-ADGB_GD or FLAG-ADGB_GD-GFP (with GFP at the C-terminal of the globin domain instead of the N-terminal) and V5-CALM in HEK293T cells. Recombinant proteins were transiently expressed and cell lysates were then incubated with anti-rabbit IgG (negative control) or anti-GFP antibody. The immunoprecipitates were immunoblotted against GFP and V5 with 2% cell lysates as input. The presence of C-terminal GFP interfere with the binding of V5-CALM onto the ADGB_GD, whereas the absence of GFP protein at the C-terminal of ADGB globin domain allows the interaction with V5-CALM, as indicated in the immunoblot against V5.

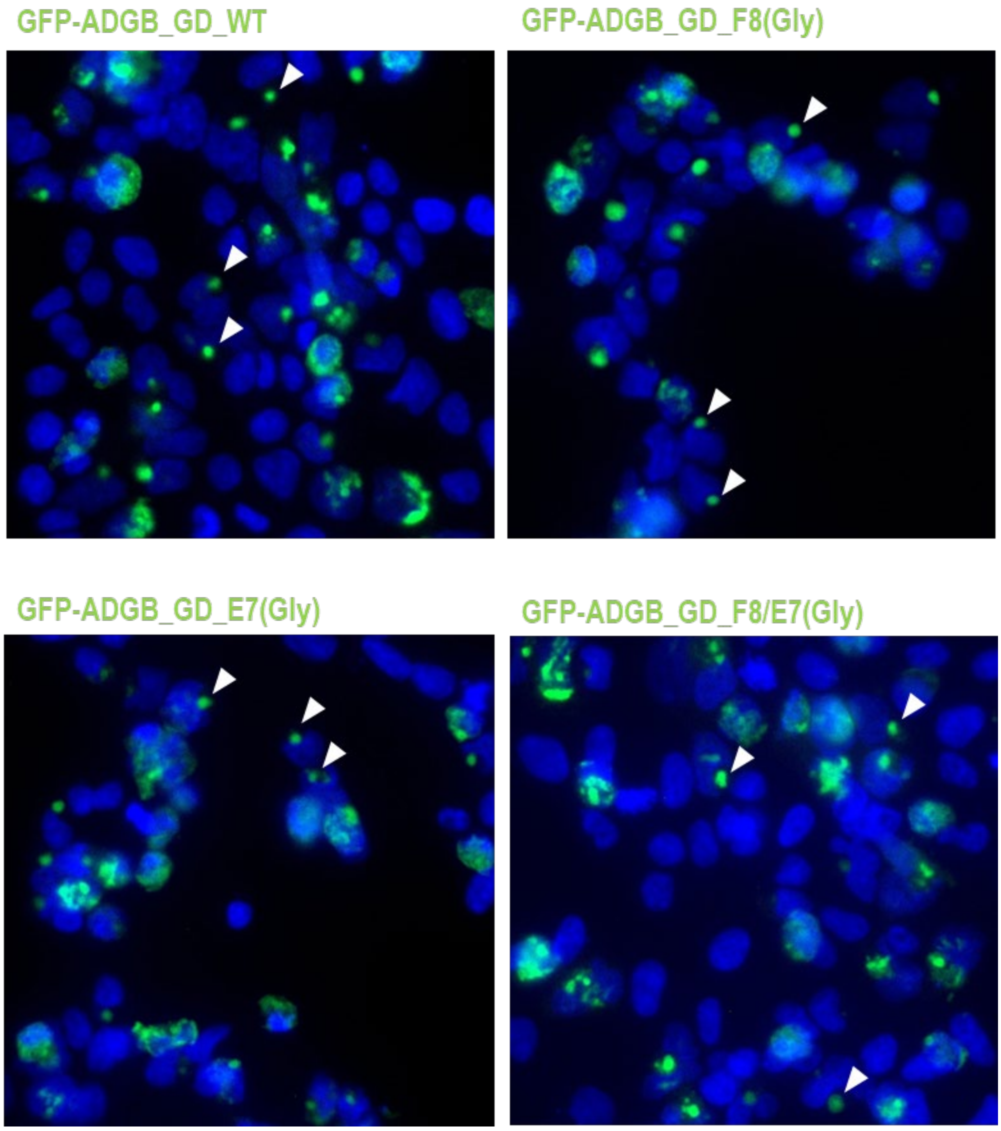


**Supplementary figure 7. Mutations on heme-binding proximal F8 and distal E7 residues within the ADGB globin domain display no change in the preferential localization to the centrosome.** FLAG-GFP-ADGB_GD-WT or FLAG-GFP-ADGB_GD-F8(Gly) or GFP-ADGB_GD-E8(Gly) or FLAG-GFP-ADGB_GD-F8/E7(Gly) were transiently expressed in HEK293T cells for 24 hours prior to 3.7% PFA/PBS fixation and mount with DAPI staining. GFP signal was observed under fluorescent microscope. Mutations on these residues show no effect on the localization of ADGB globin domain to the centrosome (PCM-1 staining in Figure 8A). Exemplary localization of signals at the centrosomes are indicated with white triangle.
